# Modular multilevel TMS device with wide output range and ultrabrief pulse capability for sound reduction

**DOI:** 10.1101/2021.09.08.459501

**Authors:** Zhiyong Zeng, Lari M. Koponen, Rena Hamdan, Zhongxi Li, Stefan M. Goetz, Angel V. Peterchev

## Abstract

**Objective:** This article presents a novel transcranial magnetic stimulation (TMS) pulse generator with a wide range of pulse shape, amplitude, and width.

**Approach:** Based on a modular multilevel TMS (MM-TMS) topology we had proposed previously, we realized the first such device operating at full TMS energy levels. It consists of ten cascaded H-bridge modules, each implemented with insulated-gate bipolar transistors, enabling both novel high-amplitude ultrabrief pulses as well as pulses with conventional amplitude and duration. The MM-TMS device can output pulses including up to 21 voltage levels with a step size of up to 1100 V, allowing relatively flexible generation of various pulse waveforms and sequences. The circuit further allows charging the energy storage capacitor on each of the ten cascaded modules with a conventional TMS power supply.

**Main results:** The MM-TMS device can output peak coil voltages and currents of 11 kV and 10 kA, respectively, enabling suprathreshold ultrabrief pulses (> 8.25 μs active electric field phase). Further, the MM-TMS device can generate a wide range of near-rectangular monophasic and biphasic pulses, as well as more complex staircase-approximated sinusoidal, polyphasic, and amplitude-modulated pulses. At matched estimated stimulation strength, briefer pulses emit less sound, which could enable quieter TMS. Finally, the MM-TMS device can instantaneously increase or decrease the amplitude from one pulse to the next in discrete steps by adding or removing modules in series, which enables rapid pulse sequences and paired-pulse protocols with variable pulse shapes and amplitudes.

**Significance:** The MM-TMS device allows unprecedented control of the pulse characteristics which could enable novel protocols and quieter pulses.

## 1. Introduction

Transcranial magnetic stimulation (TMS) devices comprise an electromagnet coil placed on the subject’s head and a pulse generator that supplies high current pulses to the coil. The coil emits intense, brief magnetic pulses, that, in turn, induce an electric field in the brain, non-invasively stimulating cortical neurons. TMS is widely used as a tool for probing and modulating brain function in research and clinical applications [1].

Existing TMS devices, however, have several significant limitations. First, TMS pulse delivery is associated with a loud clicking sound that can be as high as 140 dB resulting from electromagnetic forces in the coil [2]. The loud noise presents a risk to the hearing of the TMS subject and the operator that necessitates the use of hearing protection [2-4]. Further, by evoking an auditory response synchronized with the electromagnetic stimulus, the sound compromises the spatial localization of the stimulation effects, impacts neuromodulation, and complicates blinding, significantly impeding both basic research and clinical applications of TMS. A contributing factor to the coil sound is the typical biphasic pulse duration of 150*‒*400 µs, which corresponds to a vibration spectral peak at 5–13 kHz due to the coil winding electromagnetic forces [2, 5]. Second, existing TMS devices have limited adjustability of their pulse characteristics [6, 7]. Standard TMS devices generate sinusoidal pulses with a fixed shape and duration. More advanced devices allow some adjustment of the pulse duration and shape [8-11]. Such devices have enabled important findings regarding the effect of pulse shape and duration on neural activation thresholds [12], differential neural recruitment in the brain [13-18], lasting neuromodulatory effects [19, 20], as well as the sensation of scalp stimulation [21]. However, these devices still have a restricted range of the shape (e.g., only sinusoidal or only rectangular), duration (e.g., lacking very brief and very long pulses), and amplitude (e.g., insufficient amplitude for suprathreshold brief or complex pulses) [5, 7, 22]. Third, TMS devices have significant limitations in their ability to deliver rapid sequences of pulses for applications such as paired-pulse, triple-pulse, or quadripulse paradigms, requiring the combination of a corresponding number of individual TMS devices, and, moreover, the shape of such pulses is typically restricted to monophasic sinusoidal [23-25].

Addressing this need, we present MM-TMS, the first TMS device to use a modular multilevel circuit topology at full TMS energy levels, allowing unprecedented control of the pulse shape, width, and amplitude. This development builds upon our prior work demonstrating flexible pulse synthesis with a modular multilevel topology at lower energy levels [26].

The MM-TMS device consists of ten cascaded H-bridge modules, whose output voltages add up to form the stimulation coil voltage. This summation of the module voltages allows the generation of a significantly higher voltage (< 11 kV) than conventional TMS devices (< 2.8 kV). The available high voltages enable the generation of ultrabrief stimulation pulses (e.g., 33 µs biphasic) whose dominant vibration frequency is shifted to frequencies above the upper limit of human hearing, expected to make TMS quieter [5, 22]. Conversely, firing the modules sequentially produces very long pulses (> 400 µs), matching or exceeding the duration of conventional TMS pulses, enabling, for example, the measurement of extended strength–duration curves [12, 27].

Further, the modular topology provides 21 voltage levels within a pulse, allowing the generation of complex waveforms such as polyphasic (multicycle) pulses that can reduce the neural activation threshold and produce more robust motor evoked responses [28, 29]. The polyphasic pulses can also be amplitude-modulated, for example, with a Gaussian envelope, which has a narrower frequency bandwidth and was suggested to reduce the acoustic spectral sidebands and therefore the audible acoustic noise if the dominant frequency is concurrently shifted above the upper hearing limit [22]. Continuous kilohertz waveforms may have important neuromodulatory effects as well [30-34]. Generally, the control of the pulse characteristics over a wide range of durations and shapes may enable more selective neural recruitment and stronger neuromodulation [19, 20].

Finally, the MM-TMS device can generate rapid sequences of pulses. For example, various paired-pulse or triple-pulse paradigms [23, 24] can be implemented by instantaneously increasing or decreasing the output amplitude from one pulse to the next with the addition or subtraction of modules connected in series. Similarly, quadripulse bursts [25, 35] can be implemented by firing groups of modules in a sequence. In these pulse sequences, the parameters of the individual pulses can be controlled independently of the other pulses, although the sequential firing of module groups limits the number of voltage levels within each pulse and hence the flexibility of pulse shaping.

This paper presents the design, implementation, and electrical and acoustic characterization of an MM-TMS prototype. The measurements demonstrate the ability of the MM-TMS device to control the pulse characteristics over a wide range of shapes, widths, and amplitudes and to leverage this control for the reduction of the acoustic emissions of the coil.

## 2. Device design

### 2.1. Circuit topology

Figure 1 shows the MM-TMS circuit topology. Arm A and arm B, each consisting of five cascaded modules in our implementation, differentially drive the stimulation coil L. The coil voltage, *V*_*L*_, is the difference of the two arm voltages, *V*_AO_ and *V*_BO_. The voltage of each arm amounts to the sum of its five modules’ output voltages, where *V*_*i*_ denotes the output voltage of module *i*. Thus, the instant coil voltage is the sum of all module voltages,

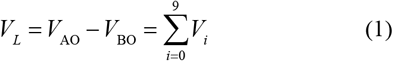

**Figure 1.**
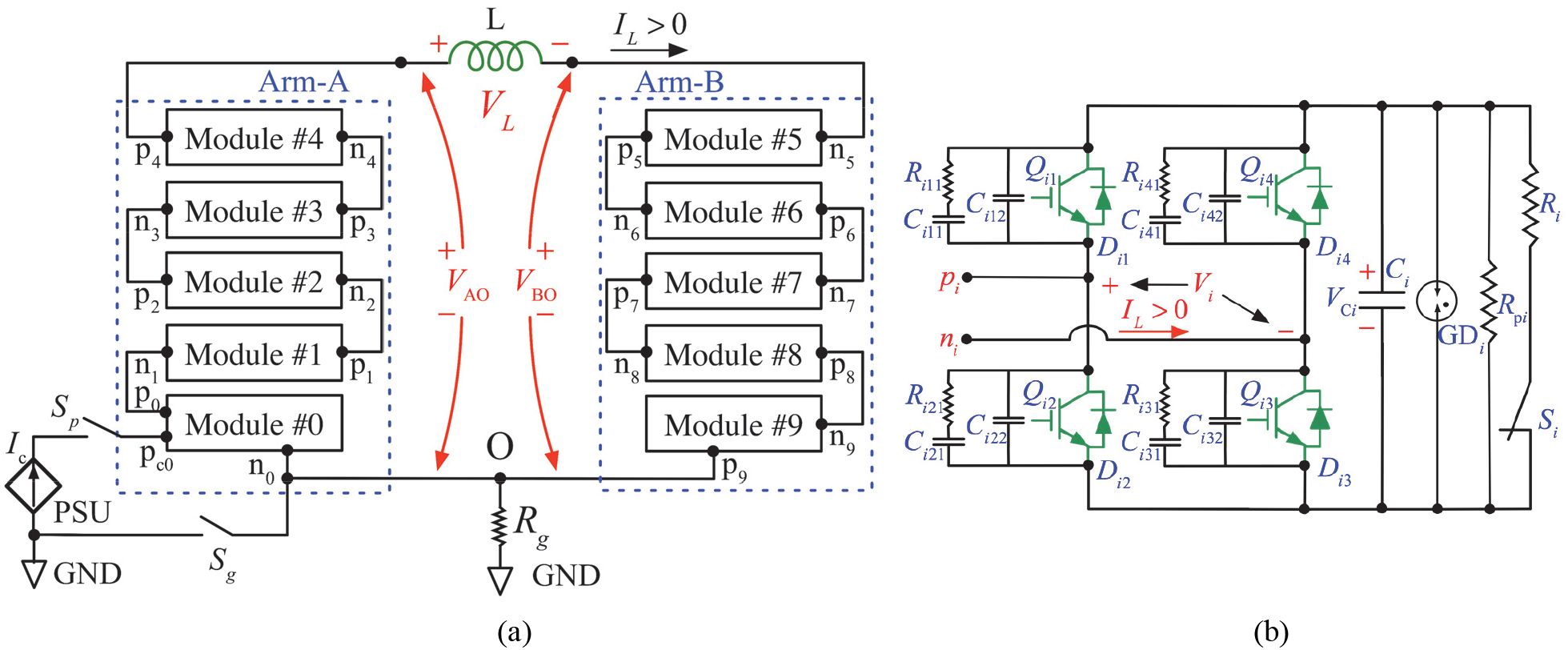
(a) Schematic diagram of the MMC-TMS pulse generator, where Arm-A and Arm-B drive the two terminals of the stimulation coil *L*, respectively, and each arm consists of five cascaded modules. The common point O of the two arms is grounded via a resistor *R*_g_. In each arm, terminal p_*i*_ of module #*i* is connected to terminal n_*i*+1_ of module #(*i*+1), where *i* = 0, 1, 2, …, 9 denotes the individual module. (b) Each module employs a full-bridge circuit interfacing an energy storage capacitor *C*_*i*_. Switches *Q*_*i*1_/*D*_*i*1_, *Q*_*i*2_/*D*_*i*2_, *Q*_*i*3_/*D*_*i*3,_ and *Q*_*i*4_/*D*_*i*4_ are IGBT modules with freewheeling diodes. Capacitors *C*_*i*12_–*C*_*i*42_ and *C*_*i*11_– *C*_*i*41,_, and resistors *R*_*i*11_–*R*_*i*41_ form snubbers. The energy storage capacitor *C*_*i*_ is discharged by a discharge resistor *R*_*i*_ when relay *S*_*i*_ is closed. In case of the relay’s failure, a secondary discharge resistor *R*_*pi*_ is directly connected to the capacitor *C*_i_, discharging the capacitor to a safe voltage below 15 V in 5 minutes. GD_*i*_ denotes gas discharge tubes (GDTs). The power supply unit (PSU) charges the module capacitors with current *I*_c_. The PSU’s positive terminal is connected to the positive terminal p_c0_ of *C*_*0*_, the storage capacitor of module #0, and its negative output is connected to bridge output n_0_ of module #0. Relays *S*_g_ and *S*_p_ disconnect the PSU from the power stage when the coil pulse is generated.

Compared to existing TMS devices, in which one terminal of the stimulation coil is grounded, the differential MM-TMS coil drive halves the system peak voltage relative to the ground and thus reduces high-voltage insulation requirements and enhances safety. For example, arm voltages of ±5.5 kV to ground can generate coil voltage of ±11 kV.

As shown in figure 1(b), each module employs an H-bridge circuit [36], implemented with high-voltage, high-current insulated-gate bipolar transistor (IGBT) switches and appropriate snubber and gate drive circuits, following our approach for prior, simpler TMS device designs [8-10, 29]. The module’s output voltage, *V*_*i*_, depends on the switches’ states and is equal to *V*_C*i*_, 0, or −*V*_C*i*_, where *V*_C*i*_ denotes the energy storage capacitor of module *i*. Typically, before a pulse is initiated at time *t* = *t*_0_, each module is charged to the same reference voltage, *V*_C*i*_ (*t* = *t*_0_) = *V*_*C*ref_, which is set by the controller. Inserting all available combinations of switch states into equation (1) yields 21 different coil voltage levels, *V*_L_ = {0, ±1, ±2, …, ±10} × *V*_*C*ref_. Hence, there are two ways of controlling the output voltage level. The first approach is to adjust *V*_*C*ref_, which provides a continuous voltage range, but is relatively slow since it requires charging or discharging the capacitors. The second approach is to adjust the number of modules connected in series, which provides discretized control of the output but is very fast, as it is limited only by the switching speed of the transistors (∼ 1 µs), thus allowing unprecedented instantaneous control over the pulse shape.

Table 1 shows the definitions of the module states with the corresponding transistor states and module output voltages. States **1**–**4** actively define the output voltage, which is determined by the commanded transistor states, regardless of the load condition. These states are typically used during all pulse phases except for the last one, allowing the pulse waveform to be accurately controlled [9-11]. For states **0** and **5**–**8**, the module output voltage depends on the diode states, which in turn depend on the circuit voltages and currents applied to the diodes. These states are used in the last phase of the pulse to automatically terminate the pulse when the coil current decays to zero as well as during the short switching transitions between phases [10].

**Table 1.** Module states and corresponding switch states and module output voltage

| Module state | Transistor switch state | | | | Module output $V_i$ |
| --- | --- | --- | --- | --- | --- |
| | $Q_{i1}$ | $Q_{i2}$ | $Q_{i3}$ | $Q_{i4}$ | |
| 0 | Off | Off | Off | Off | $\{-V_{Ci}, 0, V_{Ci}\}$ |
| 1 | On | Off | On | Off | $V_{Ci}$ |
| 2 | Off | On | Off | On | $-V_{Ci}$ |
| 3 | On | Off | Off | On | 0 (bypass) |
| 4 | Off | On | On | Off | 0 (bypass) |
| 5 | Off | On | Off | Off | $\{-V_{Ci}, 0\}$ |
| 6 | Off | Off | On | Off | $\{0, V_{Ci}\}$ |
| 7 | On | Off | Off | Off | $\{0, V_{Ci}\}$ |
| 8 | Off | Off | Off | On | $\{-V_{Ci}, 0\}$ |

### 2.2. Energy storage capacitor charging scheme

There are several options for charging the module capacitors. One approach is to embed a charger into each module. Embedded chargers, however, require ten galvanically-isolated chargers and a very high isolation voltage of ±5 max (*V*_*C*ref_) (±5.5 kV in our implementation) for the power and control signal connections to the charger. Alternatively, each of the coil terminals could be connected to a conventional grounded charger, and the modules would be charged one at a time by putting the remaining arm modules in bypass mode **[26]**. This approach requires the charger output to withstand the high voltage of ±5 max (*V*_*C*ref_) applied to the coil terminals during pulsing, or, alternatively, it requires a high voltage switch to be interposed between the charger and the coil terminals.

To circumvent the need for high-voltage isolation or switches in these approaches, we implemented a capacitor-charging scheme that requires switches with only > max(*V*_*C*ref_) voltage rating. As shown in figure 1(a), the charger’s positive output is connected to the energy storage capacitor’s positive terminal p_c0_ in module #0, and the grounded charger’s negative output is connected to output n_0_ of module #0. Figure 2 illustrates the ten modules’ state arrangements for capacitor charging. For module #0 charging *t* ∈ (*t*_1_, *t*_2_], it is in state **1** and the other modules are in state **0**. When the charger is activated (ctr is logic low), the charging current *I*_c_ goes through energy storage capacitor *C*_0_ and diode *D*_03_ of module #0. For charging of module #1 during *t* ∈ (*t*_3_, *t*_4_], *Q*_01_ of module #0 is turned on (state **7**), and modules #2 – #9 are in the bypass mode (state **3**, see Table 1), parallelizing the charger and the energy storage capacitor *C*_1_ of module #1. When the charger is activated, the charging current *I*_c_ goes through the energy storage capacitor *C*_1_ of module #1 and *D*_11_ and *D*_13_.

**Figure 2.**
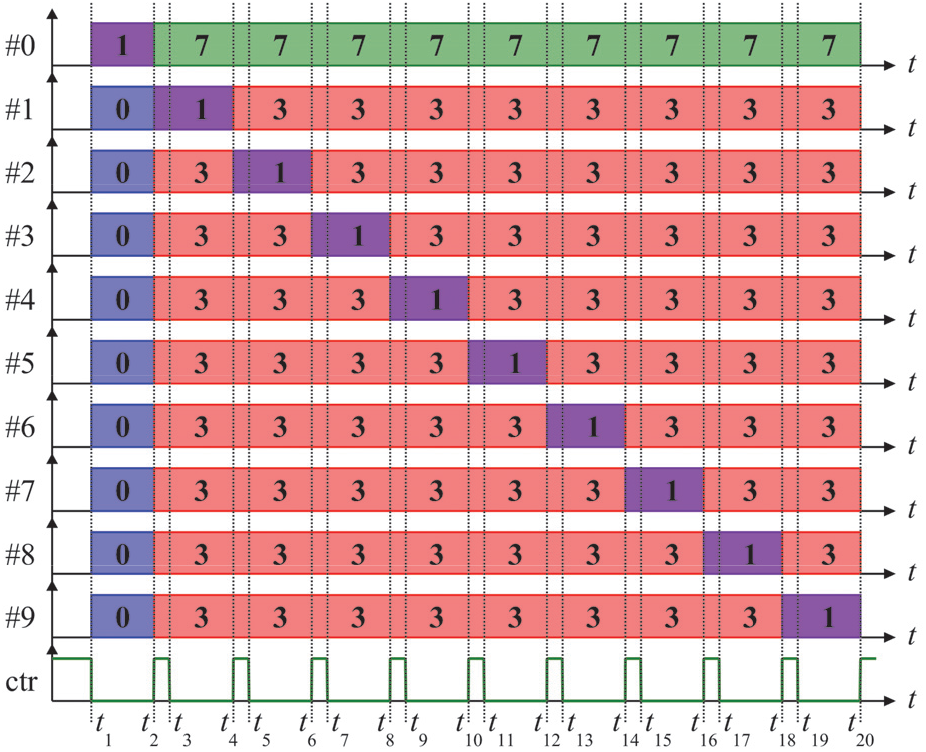
Sequence of modules states used for capacitor charging. The module capacitors are charged one at a time by the PSU. When control signal ‘ctr’ is clear, the PSU is activated, charging the module in state **1**; when ctr is set to logic high, the charging stops.

When charging modules #1 – #9, the charging current flows through the coil, but that current is very small (< 4 A). There is also a transient current between the snubber capacitors connected to node n_0_ in module #0 and the energy storage capacitor of the module being charged, which also flows through the coil. These current spikes reach 206 µs in duration but are only < 350 A in amplitude (less than 4% of the MM-TMS peak current), and therefore do not induce a significant electric field pulse (supplementary figure S12). Moreover, since the amplitude of these spikes depends on the difference between the voltages on the module #0 snubbers and the capacitor on the module being charged, the spike amplitude can be reduced to arbitrarily small values by charging the modules in several rounds by small voltage increments.

### 2.3. Pulse shape control

#### 2.3.1. Near-rectangular electric field pulses

Figure 3 illustrates four example control sequences for biphasic magnetic pulses with a nearly rectangular electric field waveform and a broad pulse width range. In figure 3(a), the ten modules’ states are synchronized. During the first pulse phase, *t* ∈ (*t*_0_, *t*_1_], each module operates in state **1**, ramping up the stimulation coil with a positive voltage of *V*_L_ = +10 *V*_Cref_. At the beginning of the second phase at *t* = *t*_1_, all modules switch to state **2**, ramping down the coil current with a negative voltage of *V*_L_ = −10 *V*_Cref_. In the third phase, starting at *t* = *t*_3_, all modules switch back to state **1**, followed by a snubbing sequence (see next section). These pulses are analogous to the biphasic pulses generated by TMS devices practically comprising a single module [9-11], but afford significantly higher output voltages, enabling briefer suprathreshold stimulation pulses.

**Figure 3.**
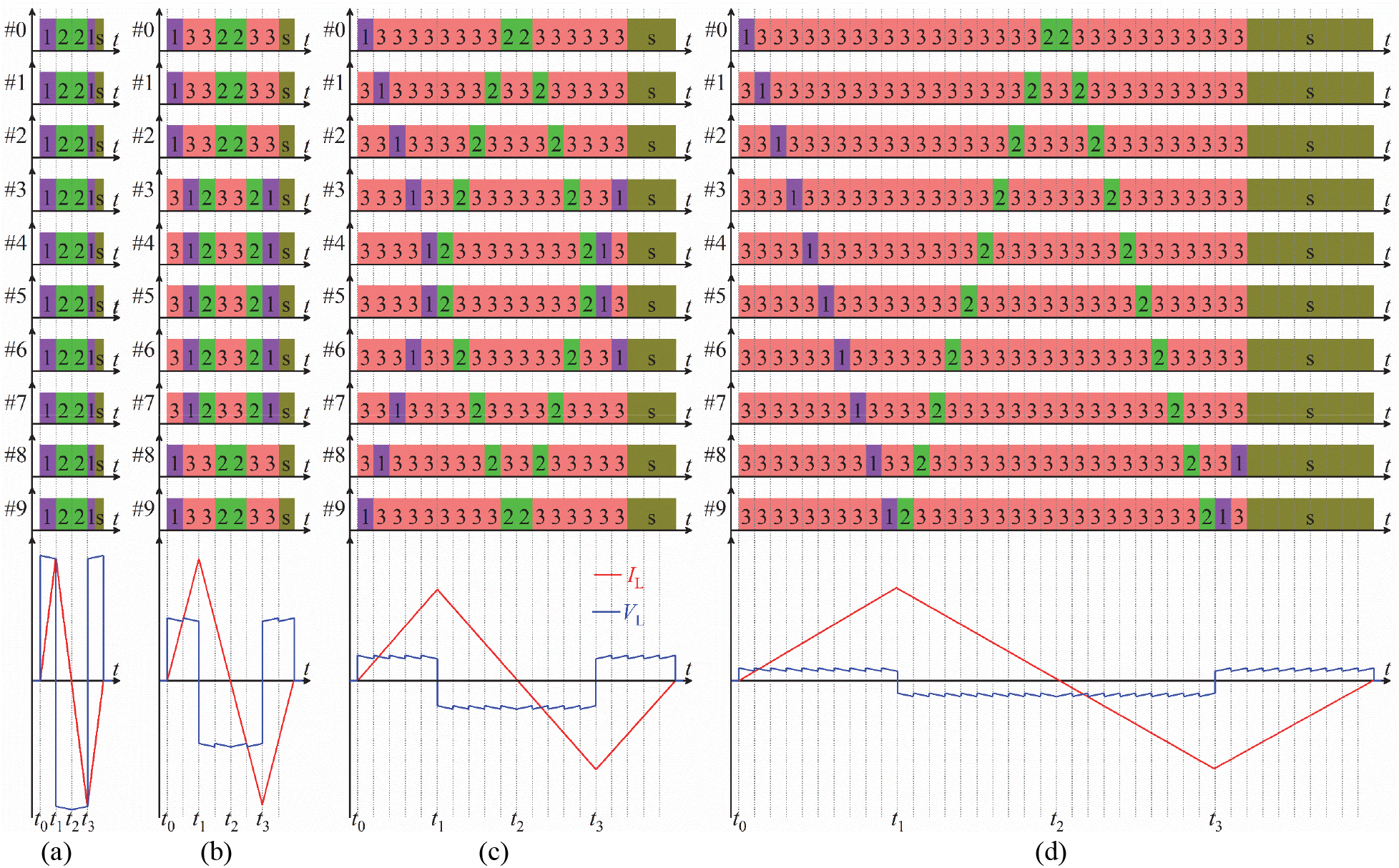
Module switching sequences for positive biphasic pulses with a wide range of pulse widths. Y-axes indicate module number, x-axes mark time intervals of the states, and colored numbers indicate switch states. State “s” denotes state sequences for active subbing, which does not affect the circuit behavior until the coil current decays to zero at the end of the TMS pulse, when the snubbing starts to dampen the coil ringing (see Supplementary Material). (a) For the briefest pulses, the ten modules’ switch states are synchronized. (b) Two module groups consisting of five modules in series connection sequentially drive the coil (5+5 scheme). (c) Five module groups consisting of two modules in series connection sequentially drive the coil (2+2+2+2+2 scheme). (d) Each of the ten modules sequentially drives the coil (1+1+1+1+1+1+1+1+1+1 scheme). Experimental implementations of these pulses are shown in figure 6.

However, the synchronized scheme in figure 3(a) can only be used for relatively brief pulses because of the small total series capacitance of *C*_*i*_/10 driving the coil. To generate longer pulses, the control schemes in figures 3(b)–(d) fire groups of modules sequentially. Sequential firing reduces the available peak coil voltage, but this is acceptable since longer electric field pulses require lower amplitudes for suprathreshold stimulation.

In figure 3(b), the ten modules are divided into two groups, each of which consists of five modules, and in each module group, the modules’ states are synchronized. For example, modules #0, #1, #2, #8, and #9 form a module group, and the remaining modules (#3 – #7) form the other group. We refer to this scheme as ‘5+5’. During the pulse, one module group is activated at a time, while the other is in a bypass state, imposing ±5*V*_*C*ref_ on the stimulation coil. For example, when *t* ∈ (0, *t*_1_], the two module groups sequentially charge the stimulation coil with a voltage of +5*V*_*C*ref_.

In figure 3(c), two modules, both of which have the same state, form a group. In this ‘2+2+2+2+2’ scheme, the five groups are formed by modules {#0, #9}, {#1, #8}, {#2, #7}, {#3, #6}, and {#4, #5}, respectively. At any instant during the pulse, one module group charges or discharges the coil with the remaining four module groups in the bypass state, leading to a coil voltage of ±2*V*_*C*ref_. For example, during *t* ∈ (0, *t*_1_], the five module groups sequentially charge the stimulation coil with a voltage of +2*V*_*C*ref_.

In figure 3(d), each module sequentially charges or discharges the stimulation coil with the remaining nine modules in the bypass state, imposing ±*V*_*C*ref_ on the stimulation coil. In this ‘1+1+1+1+1+1+1+1+1+1’ scheme, the ten modules sequentially charge the stimulation coil with a voltage of +*V*_*C*ref_ during *t* ∈ (0, *t*_1_].

The control schemes have important symmetries. First, at each time during a pulse, the device uses an equal or approximately equal number of modules from the two power stage arms (A and B). The symmetric involvement of both arms ensures that the coil common-mode voltage is close to zero [26, 37]. Second, the module groups’ state timings are symmetric with respect to the phase transition points to avoid severe unbalance of the capacitor voltages due to different discharging and charging currents. For example, during *t* ∈ (0, *t*_2_], the modules’ state timing is symmetric with respect to *t* = *t*_1_, ensuring approximately the same discharging and charging current for each module during a pulse. Since the discharging and charging of the capacitors is largely symmetric, most of the pulse energy is returned to the capacitors at the end of the pulse, and this energy can be recycled from pulse to pulse, as in other efficient repetitive TMS devices [6, 9-11, 38]. Note that schemes with +3*V*_*C*ref_ and +4*V*_*C*ref_ are also possible in ‘3+3+3’ and ‘4+4’ schemes by setting the unused 1 or 2 modules, respectively, to bypass mode for the duration of the whole pulse.

Using the same approach, we can also develop module state timings for monophasic magnetic pulses with a wide range of pulse widths (a monophasic pulse is essentially half of a biphasic pulse). Electric field pulses with asymmetric phase amplitude and duration [9, 10] can be generated by combining brief, high-amplitude phases of synchronous firing of the modules with long, low-amplitude phases of sequential firing of the modules. Inversion of the pulse voltage polarity is trivially achieved in MM-TMS by flipping the polarity of the output voltage of each module.

Finally, schemes using sequential firing of the modules should consider limitations on the maximum current that the IGBTs can withstand during the hard (forced) commutation when a module switches from diode conduction to IGBT conduction of the current [39].

#### 2.3.2. Complex pulse waveforms

The multilevel topology of MM-TMS enables the generation of complex pulse waveforms. For example, the device can approximate a sinusoidal polyphasic coil voltage *V*_L_ and resultant coil current *I*_L_ with a Gaussian amplitude envelope by staircase discretization of the waveform. Due to the device’s rich redundancies of module states, various modulation schemes can produce such a waveform. Considering the trade-off between the practicality and approximation accuracy, we designed a modulation scheme consisting of 61 modulation states, where each state has the same duration. As illustrated by figure 4, two modules construct a module group, and the module states are synchronized within each of the five module groups. The modulation scheme approximates a sinusoidal polyphasic coil voltage with a Gaussian amplitude envelope by connecting different module groups in series. Using redundant module states, the modulation scheme balances the module voltages by discharging and charging the same capacitor with approximately the same coil current and duration.

**Figure 4.**
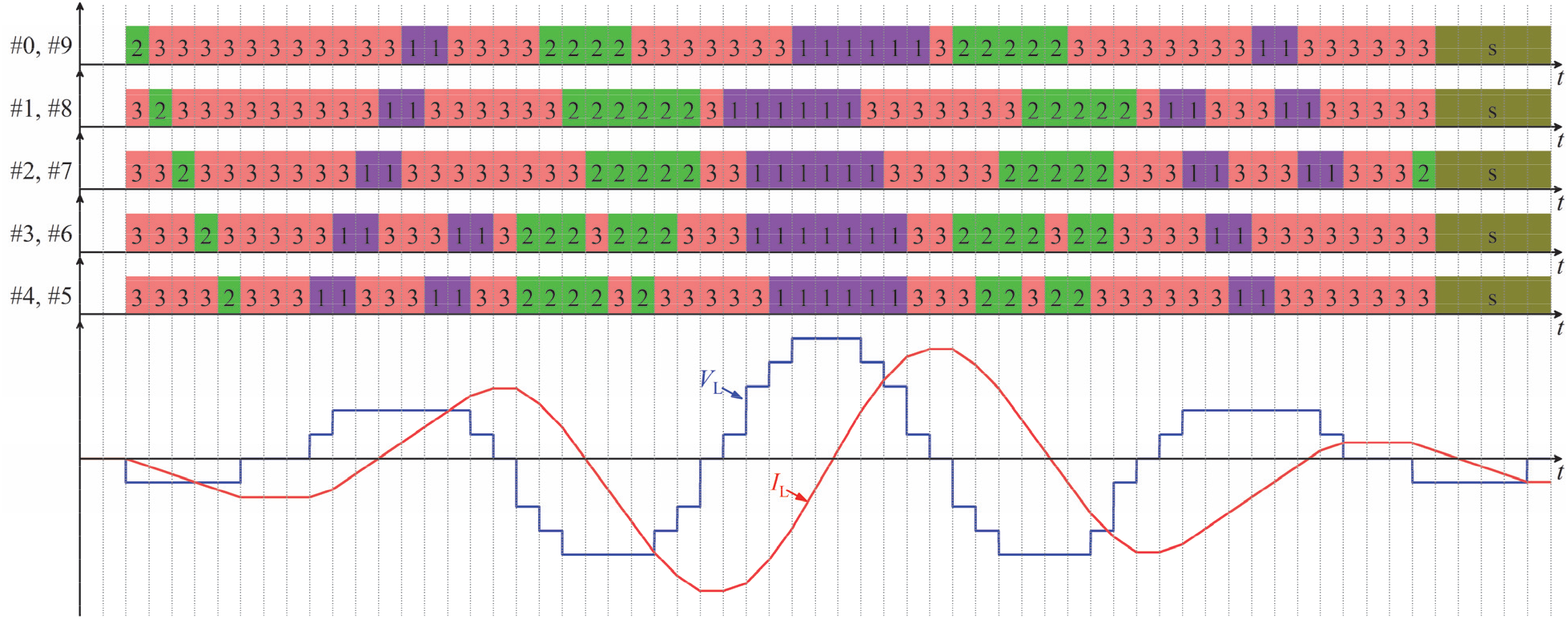
Module state sequence (top) to generate a polyphasic sinusoidal pulse with a Gaussian amplitude envelope (bottom). State diagram conventions are as in figure 3. For simplicity the illustration is for constant module capacitor voltages.

### 2.4. Pulse snubbing

For the pulse application featuring large currents, the snubbers are essential to limit the potentially damaging voltages and currents in semiconductors during switching. Figure 1(b) shows the snubbers included in each module of the proposed topology. During IGBT turn-off, the capacitors *C*_*i*12_, *C*_*i*22_, *C*_*i*32,_ and *C*_*i*42_ primarily serve to take over a portion of the IGBT current, while *C*_*i*11_–*R*_*i*11_, *C*_*i*21_–*R*_*i*21_, *C*_*i*31_–*R*_*i*31_, and *C*_*i*41_–*R*_*i*41_ mainly dampen the voltage ringing across the collector and emitter during turn-off transients [9, 10]. The IGBT’s minimum snubbing requirement determines the capacitance of *C*_*i*12_, *C*_*i*22_, *C*_*i*32_, and *C*_*i*42_ since a larger capacitance increases the switching loss and stresses the IGBT during turn-on.

The snubber capacitors, however, cause ringing at the end of each pulse [9], [10]. Therefore, we deployed active snubbing, which uses the module IGBTs to dissipate the snubber capacitor energy at the end of a pulse. We extended our prior active snubbing approach [10] by leveraging the multi-module topology of MM-TMS to interleave the snubbing switching across the modules, which enabled smoother damping waveforms and reduced switching frequency of the individual IGBTs compared to snubbing with synchronized module switching. Details of the active snubbing switching sequences and performance are presented in the Supplementary Material.

In the MM-TMS device, we use low-inductance bus bar connections between the energy-storage capacitor *C*_*i*_ and switches *Q*_*i*1_*/D*_*i*1_ – *Q*_*i*2_*/D*_*i*2_ and *Q*_*i*3_*/D*_*i*3_ – *Q*_*i*4_*/D*_*i*4_. This obviates the need for snubbers across the IGBT half-bridges, which were necessary in other IGBT-based TMS devices [10].

### 2.5. Circuit implementation

We constructed a ten-module MM-TMS device based on the circuit in figure 1, with maximum coil voltage of 11 kV and maximum coil current of 10 kA, which we had estimated to be necessary for suprathreshold cortical stimulation with ultrabrief pulses (33 µs biphasic) [22]. Table 2 summarizes the parameters of the key components of the device. The device was assembled in a 35U Sound Control Cabinet (Rackmount Solutions, TX, USA) cabinet, which suppresses the emission of pulsed sound by the power components and interconnections.

**Table 2.**
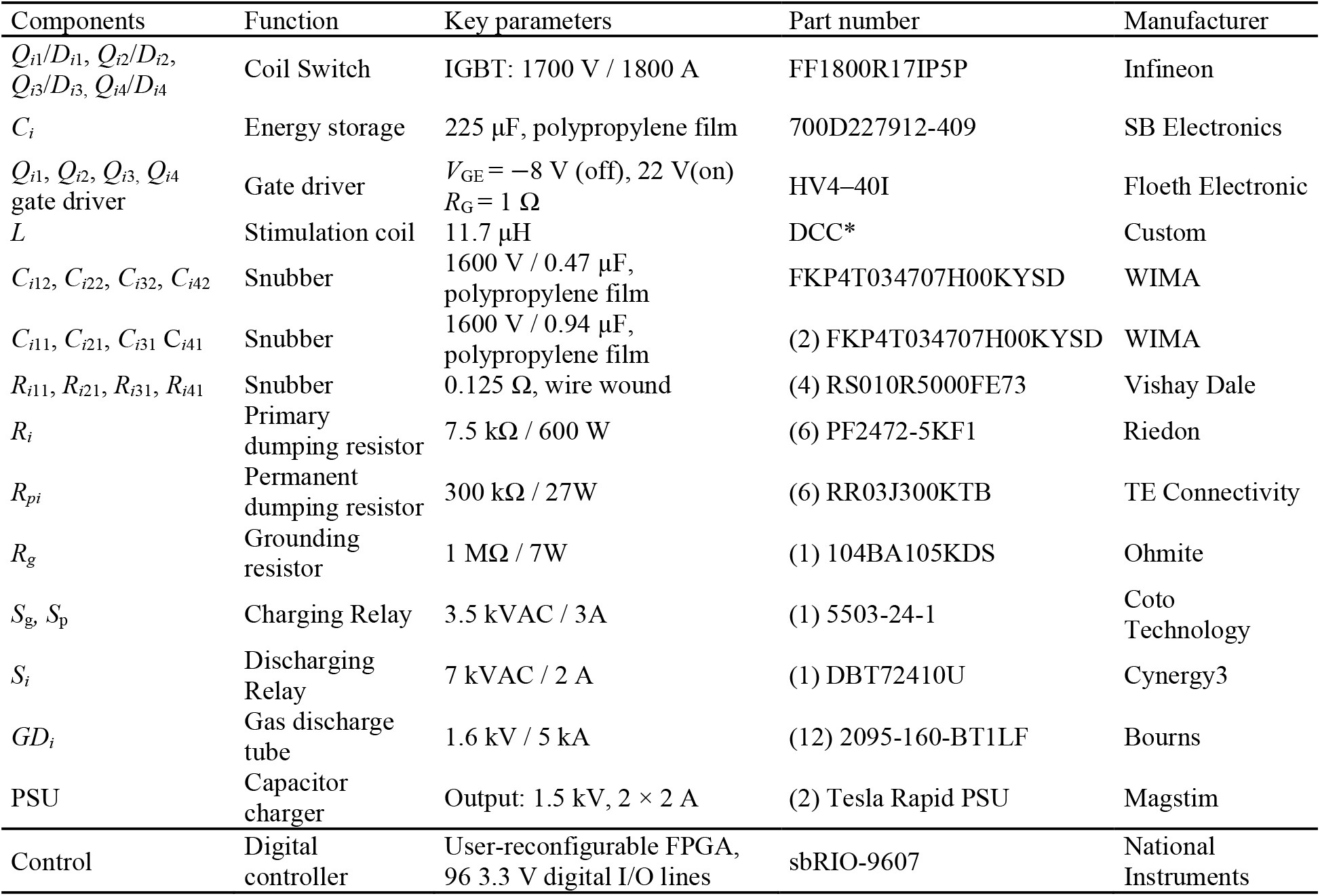
Key MM-TMS circuit implementation components.

#### 2.5.1. Energy storage capacitors and charging

For the proposed 11 kV/10 kA pulse generation, the voltage rating of the energy storage capacitor *C*_*i*_ in each module must be larger than 1.1 kV, and its peak-current rating must be larger than 10 kA. Given the inductance (11.7 µH) of the quiet stimulation coil designed for the MM-TMS device [40] (see section 2.5.5), the total capacitance seen by the coil should range from 20 µF to 35 µF to achieve not only the required minimal biphasic pulse duration of 33 µs, but also a range of longer pulse widths and efficient near-rectangular electric field pulse shapes [8, 10, 41]. Since the modules are connected in series, the energy storage capacitance of each module should therefore be 200 µF to 350 µF. In addition, to handle fast switching of large current, it is essential to implement a capacitor with a low equivalent series inductance to suppress the voltage spike during the IGBT’s turn-off. Furthermore, the low equivalent series resistance is critical to limit losses and temperature rise with the large current. Based on these considerations, we selected a power ring film capacitor for *C*_*i*_ with low stray inductance (< 5 nH) and resistance (< 250 µΩ), high voltage rating (1.2 kV dc), and extreme current for small duty ratios (> 7.5 kA). The module output is protected from a significant overvoltage by gas discharge tubes in parallel with the capacitor.

Using the charging scheme described in section 2.2, the ten modules’ energy storage capacitors are charged sequentially by two parallel Magstim power supply units (PSUs) controlled by a custom electronics interface. The capacitor voltage is sensed by a voltage divider and fed to one input of a comparator. The comparator’s other input is fed from a digital-to-analog converter programmed with the controller’s scaled target voltage, *V*_*C*ref_. When the target voltage is reached, the comparator’s output is set to logic high, and the charging stops.

#### 2.5.2. IGBT switches and gate drivers

Switches *Q*_*i*1_*/D*_*i*1_ ‒ *Q*_*i*2_*/D*_*i*2_ and *Q*_*i*3_*/D*_*i*3_ ‒ *Q*_*i*4_*/D*_*i*4_ in each module are implemented with a half-bridge IGBT module rated at 1.7 kV and 1.8 kA (FF1800R17IP5P). The half-bridge module integrates the upper and lower IGBT (for example, *Q*_*i*1_ and *Q*_*i*2_) and their freewheeling diodes (*D*_*i*1_ and *D*_*i*2_, respectively), minimizing the parasitic inductance and suppressing the turn-off voltage spike. Given the maximum output peak voltage of 11 kV, each module has a maximum working voltage of 1.1 kV; with rating voltage of 1.7 kV, FF1800R17IP5P has therefore a 600 V safety margin to accommodate the voltage spikes introduced by stray inductance during switching transients.

FF1800R17IP5P has a dc current rating of 1.8 kA and a repetitive peak current rating of 3.6 kA for 1 ms pulses. While this is below the MM-TMS device’s maximum pulse current of 10 kA, below a junction temperature of 125 ^**o**^C, IGBTs can withstand a brief current that is about ten times larger than its rating current [42-44]. The junction temperature depends on the switching loss that consists of the conducting losses and switching losses. As discussed in section 2.4, snubbers are added to partly take over the current during switching, reducing the switching loss. Further, it is necessary to reduce the conducting loss by decreasing the IGBTs’ on-state voltage and preventing desaturation at high currents. Therefore, we shifted the voltage range of a commercial gate driver to produce a gate–emitter voltage of 22 V in the on-state, which is higher than the standard value of 15 V. The selected gate voltage is an improved trade-off between the IGBTs’ on-state voltage and gate–emitter voltage limit. Besides, the gate driver’s output resistance is set to the minimum value (R_G_ = 1 Ω) determined by the gate driver’s maximum output current, ensuring a switching time of about 1 µs.

#### 2.5.3. Connections within and between modules

For the proposed MM-TMS device featuring fast switching with a large current, it is essential to minimize any stray inductance to optimize the performance, including the stray inductance of the connection between IGBTs and the energy storage capacitor on each module as well as the interconnections between modules. Minimized parasitic inductance of the connection between the IGBTs and the energy storage capacitor suppresses the IGBT voltage spike during switching, whereas minimized parasitic inductance of the connection between the modules increases energy transfer to the stimulation coil. Therefore, to minimize stray inductance, these connections are implemented with laminated bus bars that consist of two copper plates separated by a thin dielectric material and laminated between insulating sheets. For example, in figure 1(a), *n*_9_ and *p*_8_ are connected by one layer of the laminated bus bar, and the other layer connects *p*_0_ and *n*_1_. Since these two module interconnections both carry the coil current but in opposite directions, the stray magnetic flux is largely cancelled.

#### 2.5.4. Snubbers

The snubbers of each IGBT half-bridge are implemented with a two-layer printed circuit board (PCB), which is mounted on the terminal connectors of the laminated bus bars mounted on the IGBT modules. The components on the PCB were laid out to optimize the snubber’s performance. Snubber capacitors *C*_*i*12_, *C*_*i*22_, *C*_*i*32_, and *C*_*i*42_ were placed close to the IGBT terminals, minimizing the parasitic inductance. The snubber component values, listed in Table 2, were determined by the minimum requirements on IGBT turn-off current take-over and voltage-spike suppression.

#### 2.5.5. Stimulation coil

The MM-TMS device is connected to a double containment coil (DCC*) which we developed previously [40]. This coil was optimized for reduced acoustic noise and ultrabrief pulses, using litz wire windings for low high-frequency losses. The coil is designed for the internal differential working voltage of 11 kV and to provide two means of patient protection for the ground-referenced working voltage of 5.5 kV. The coil is connected to the MM-TMS device with a 5 m power cable terminated with compression lugs bolted directly to the laminated bus bar connecting the modules. The omission of a coil connector largely compensates for the longer power cable, resulting in the total coil inductance (11.7 µH) and resistance (30 mΩ) closely matched to standard TMS coils.

#### 2.5.6. Controller

The MM-TMS device is controlled by a single-board reconfigurable I/O compact controller (sbRIO-9607, National Instruments, USA), which integrates a real-time processor, a user-reconfigurable field-programmable gate array (FPGA), and 96 3.3 V digital I/O lines. With a 40 MHz oscillator, the FPGA provides precise timing with a 25 ns resolution for the 96 digital I/O lines. The custom electronics interface between the power circuits and the sbRIO-9607 implements capacitor voltage sensing, charging control, coil current and temperature sensing, IGBT gate driver interface, as well as other device control and monitoring functions ensuring safe operation. The sbRIO-9607 is controlled remotely by a host computer running a LabVIEW (National Instruments) graphical user interface, where the user can specify the TMS pulse parameters.

## 3. Experimental methods

### 3.1. Electrical measurements

The stimulation coil voltage was obtained with two differential voltage probes measuring arm voltages *V*_AO_ and *V*_BO_, respectively. The coil current was measure with a commercial Rogowski current sensor. The electric field, *E*, was measured with a PCB-based single-turn search coil fixed on the MM-TMS coil [2]. The measurements were recorded with a digitizing oscilloscope.

### 3.2. Stimulation strength estimation

The stimulation strength of TMS pulses, i.e., their ability to depolarize cortical neurons, depends on the pulse shape, duration, and amplitude. Therefore, all of these factors have to be considered when comparing different pulse types. The depolarization of cortical neuron membranes by the TMS pulses was estimated with a linear first-order low-pass filter, where the neural membrane voltage change, Δ*V*_m_, is the filtered output of the measured electric field *E* [27]. The time constant of the low pass filter was set to 200 µs, which was estimated empirically from strength–duration curves for motor cortex activation with TMS pulses [12]. The estimated neural membrane voltages are normalized by an average resting motor threshold (RMT) to facilitate the stimulation strength quantification of various TMS pulses [2]. The first-order lowpass filter was implemented with the filter function in MATLAB (The MathWorks, USA). However, this linear model may overestimate the stimulation strength of ultrabrief rectangular pulses as it neglects ion-channel dynamics [45, 46], and underestimates the stimulation strength of polyphasic pulses [28, 29, 47]. Thus, the peak Δ*V*_m_ value should be interpreted only as a rough approximation for the effective stimulation strength.

### 3.3. Acoustic measurements and analysis

The short-duration impulsive sound produced by TMS was recorded with a set-up we described previously [2, 40]. Briefly, an omnidirectional pressure microphone (Earthworks M50, Earthworks Audio, USA) was placed 25 cm from the center of the head-facing side of the coil, amplified with a wide-input-range preamplifier (RNP8380, FMR Audio, USA) and sampled with a 192 kHz audio interface (U-Phoria UMC404HD, Behringer, Germany). Then, we used the electromagnetic artifact suppression method and band-pass filters described with 0.08–50 kHz bandwidth from our previous study [2]. Finally, to separate the weak sound of ultra-brief pulses from the ambient noise present in our laboratory, we averaged 20 trigger-synchronized TMS pulses per measurement condition. Given the lower sound levels, we only measured sound for pulses at 167% RMT and at 251% RMT. The spectra of the coil sound and the pulse loudness were computed using methods described previously [40].

## 4. Experimental results

### 4.1. Wide electric field amplitude range

Figure 5 illustrates the two schemes for controlling the output electric field amplitude of MM-TMS, discussed in section 2.1. The first scheme connects the ten modules in series and progressively increases the capacitor voltage, whereas the second scheme increases the numbers of the modules in series from 1 to 10, which allows for faster, but discrete, amplitude adjustment. For a module capacitor voltage of 1000 V, the output reaches 1300 V/m, which can be increased further by 10% for the maximum designed module voltage of 1100 V. In contrast, conventional TMS devices induce peak electric field ranging approximately 125–250 V/m [48].

**Figure 5.**
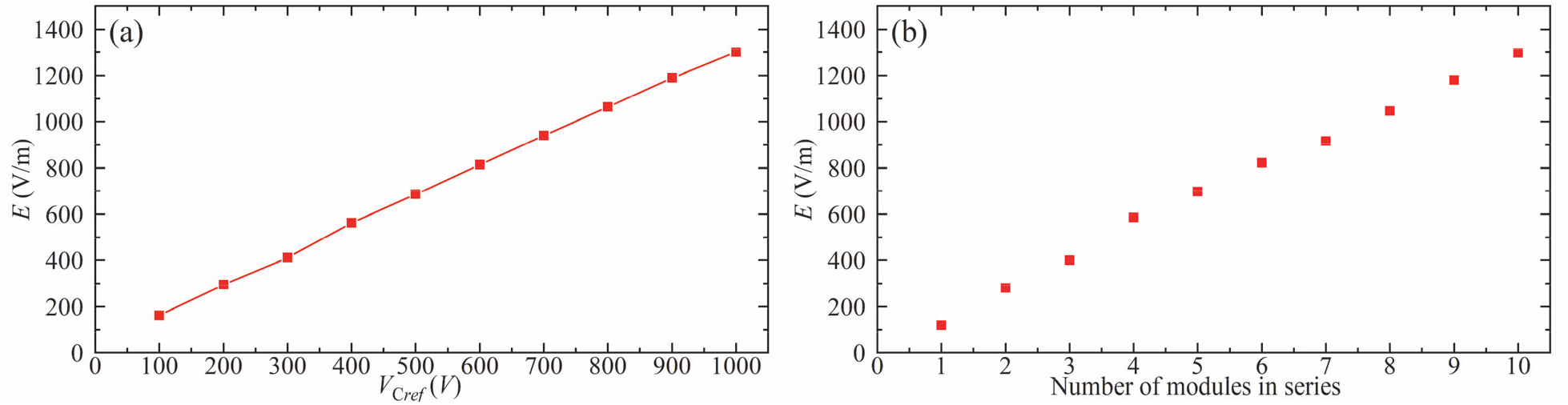
Two schemes to control the MM-TMS output electric field amplitude. Measured peak electric field, *E*, of a positive biphasic ultrabrief pulse with a pulse width of 33 μs for (a) the ten modules connected in series and with various capacitor voltages for *V*_*C*ref_ from 100 V to 1000 V and (b) different number of modules connected in series and with fixed *V*_*C*ref_ = 1000 V.

### 4.2. Wide pulse width range

Figure 6 illustrates positive biphasic pulses with a wide range of pulse widths. Synchronized module switching was used to generate brief high-voltage pulses shown in figures 6(a)–(f). The three brief pulses have the same coil voltage (10 kV) but different pulse widths (33 μs–50 μs), leading to different peak currents (7 kA–8.5 kA).

**Figure 6.**
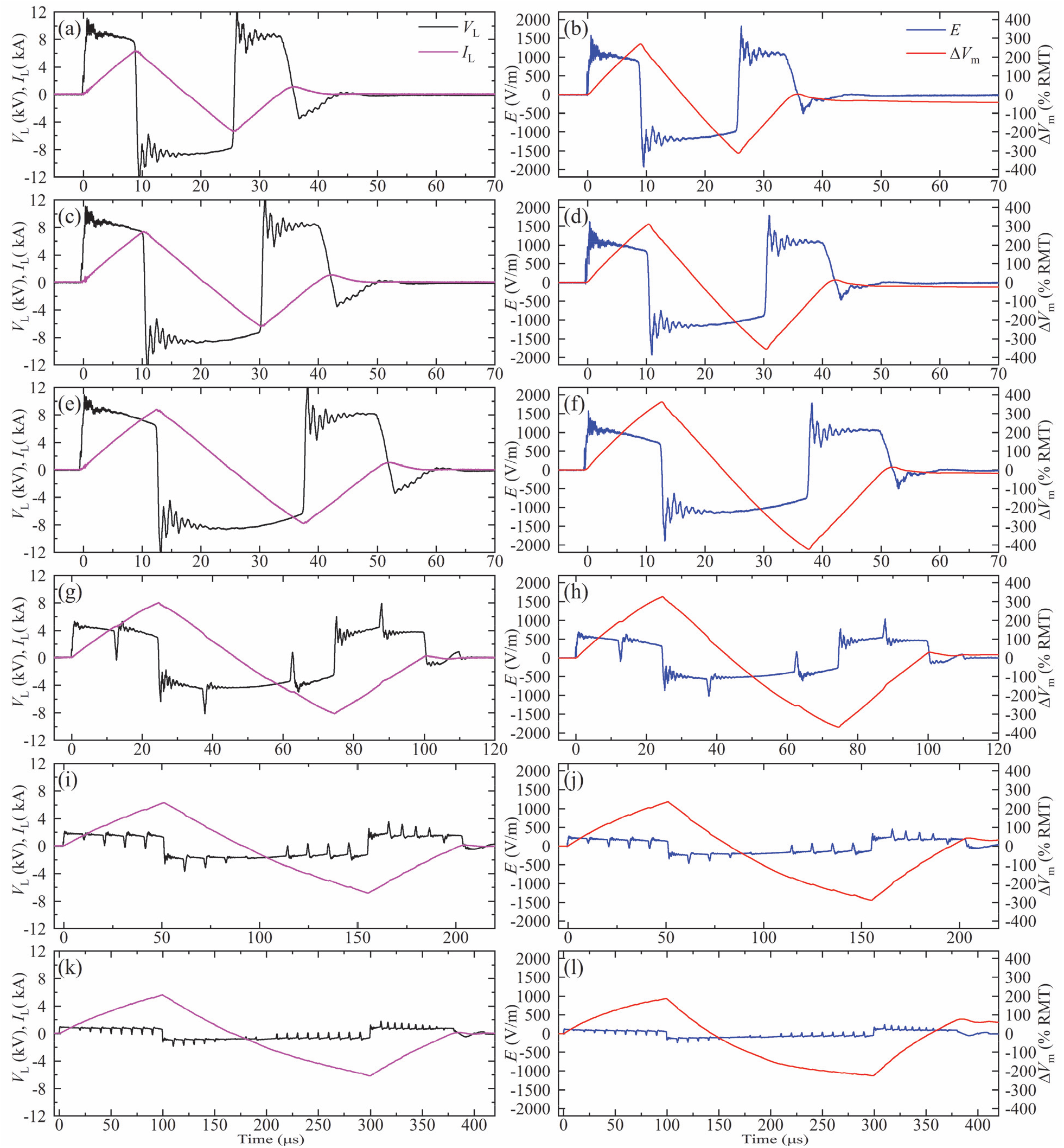
Measured coil current *I*_L_, coil voltage *V*_L_, electric field *E*, and estimated neural depolarization Δ*V*_m_ of positive biphasic pulses with *V*_*C*ref_ = 1100 V and various widths: (a), (b) 33 μs; (c), (d) 40 μs; (e), (f) 50 μs; (g), (h) 100 μs; (i), (j) 200 μs; (k), (l) 400 μs. These pulses use the module switching schemes described in figure 3: the pulses in (a)–(f) are generated with all modules connected in series, and the remaining rows correspond to the 5+5, 2+2+2+2+2, and 1+1+1+1+1+1+1+1+1+1 sequential module activation schemes, respectively.

Figures 6(g)–(h) demonstrate a 100 μs positive biphasic pulse generated with the 5+5 scheme, where the two module groups are sequentially activated for 12.5 μs to drive the coil with ±5 kV. Similarly, figures 6 (i)–(l) demonstrate 200 μs and 400 μs positive biphasic pulses using the 2+2+2+2+2 and 1+1+1+1+1+1+1+1+1+1 schemes, respectively, where each group of modules is on for 10 μs. These sequential module activation schemes enable markedly longer near-rectangular pulses than could be achieved by a single module or series connected modules.

The estimated neural membrane depolarization Δ*V*_m_ for all pulses in figure 6 exceeds 200% RMT, suggesting that these pulses would produce suprathreshold stimulation in most subjects.

In figures 6 (g)–(l), brief (< 1.5 μs) voltage and associated electric field spikes occur during the modules’ switching due to the small differences of the switching speed between the modules. Nevertheless, these spikes are too brief to significantly affect the neural membrane potential, as confirmed by the Δ*V*_m_ traces.

### 4.3. High-amplitude ultrabrief monophasic and biphasic pulses

While figure 6 illustrated only positive biphasic magnetic pulses, figure 7 demonstrates also high-amplitude ultrabrief (8.25 μs initial phase) monophasic and biphasic pulses of both polarities. This illustrates the flexibility of the MM-TMS device, including electronic pulse polarity control.

**Figure 7.**
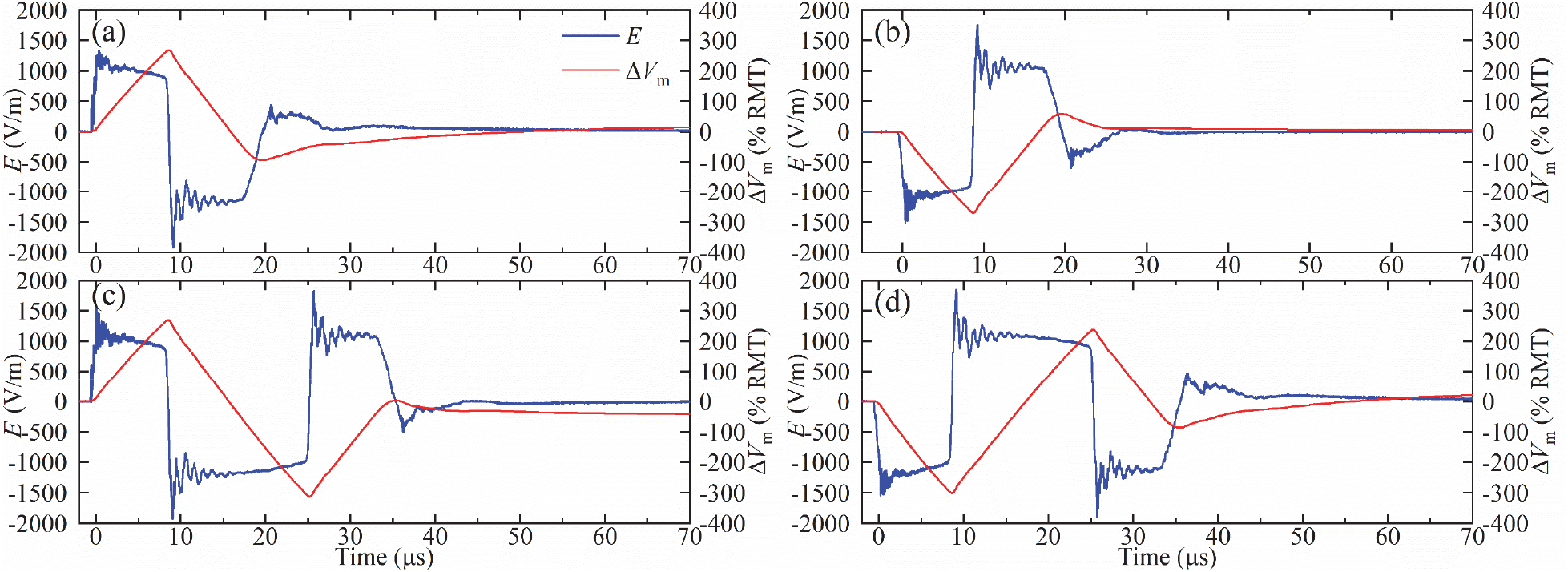
Measured electric field *E* and estimated neural depolarization Δ*V*_m_ of (a) positive and (b) negative 16.5 μs monophasic pulse, and (c) positive and (d) negative 33 μs biphasic pulse for *V*_*C*ref_ = 1100 V.

Further, by bypassing all the modules during a pulse, the prototype can insert a zero-voltage phase (interphase) with a controllable duration between the negative and positive electric field phases, which can reduce the neural activation threshold without increasing the peak coil current [49-52]. Figure 8 demonstrates the four pulses configurations from figure 7 but with interphases, where each pulse has an initial phase of 8.25 μs duration and an interphase of 8.5 μs.

**Figure 8.**
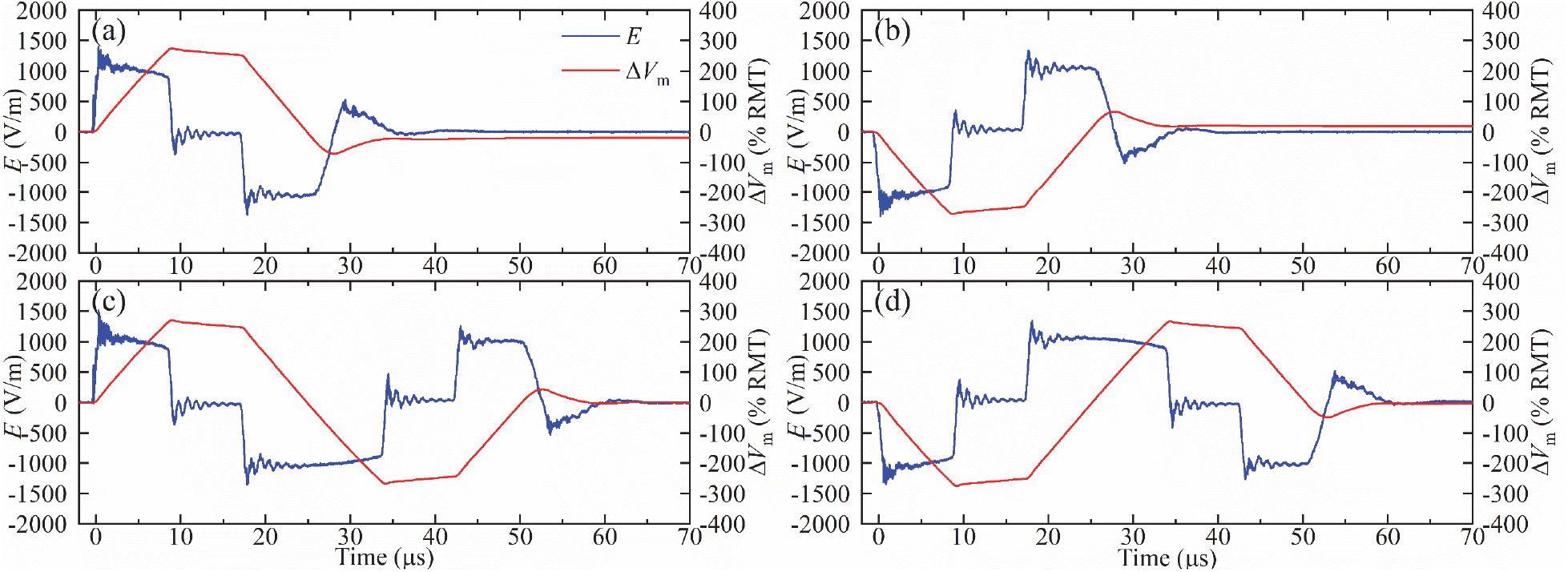
Measured electric field *E* and estimated neural depolarization Δ*V*_m_ of (a) positive and (b) negative 25 μs monophasic pulse with an 8.5 μs interphase, and (c) positive and negative (d) 50 μs biphasic pulse with 8.5 μs interphases for with *V*_*C*ref_ = 1100 V. The interphases cause the coil current to be approximately trapezoidal.

### 4.4. Short-interval paired pulses

The MM-TMS device’s unprecedented control over pulse parameters and its fundamentally energy-lossless topology allow the generation of paired-pulse protocols with various pulse shapes that conventionally require the outputs of two devices to be combined. The device can instantaneously increase the output amplitude from one pulse to the next by adding more modules in series, generating paired pulses with different stimulation strength.

Figures 9(a) and 9(b) illustrate this capability with a pair of monophasic positive pulses delivered with a short inter-stimulus interval (1 ms) with the stimulation strength increasing by 25% and 34%, respectively, for the second compared to the first pulse. This is achieved by connecting all ten modules in series for the second pulse while connecting only seven and six modules in series, respectively, for the first pulse.

**Figure 9.**
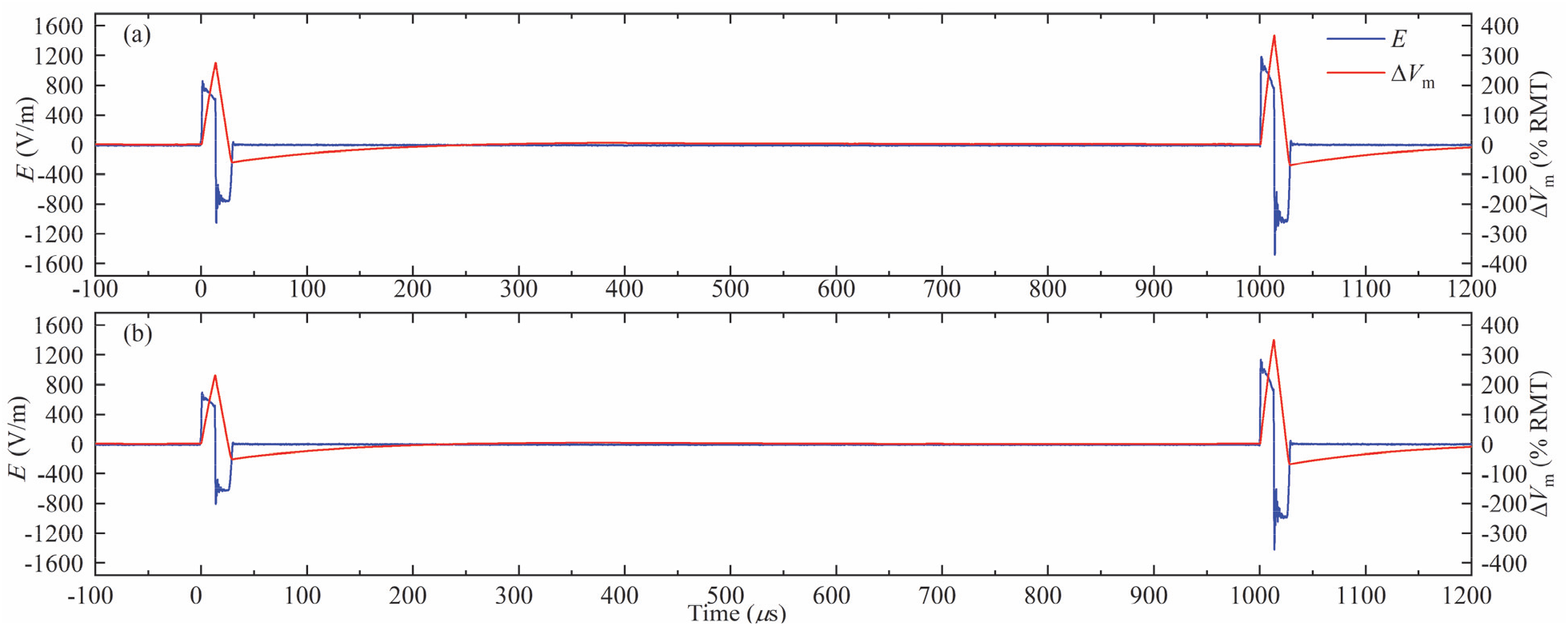
Pair of positive monophasic pulses delivered with a short interstimulus interval (1 ms). (a) The first pulse has a lower stimulation strength than the second pulse by 25%. (b) The first pulse has a lower stimulation strength than the second pulse by 36%. A longer, 3 ms interstimulus interval is illustrated in supplementary figure S11.

### 4.5. Amplitude-modulated sinusoidal polyphasic pulses

The capability of MM-TMS to generate relatively complex pulse shapes is illustrated in figure 10 with sinusoidal polyphasic pulses with a Gaussian envelope. Compared to a conventional polyphasic pulse with a flat envelope [28, 29], these pulses reduce the subharmonic sideband in the spectrum of the coil current, which could improve the acoustic performance of polyphasic pulses, as discussed in the following sections.

**Figure 10.**
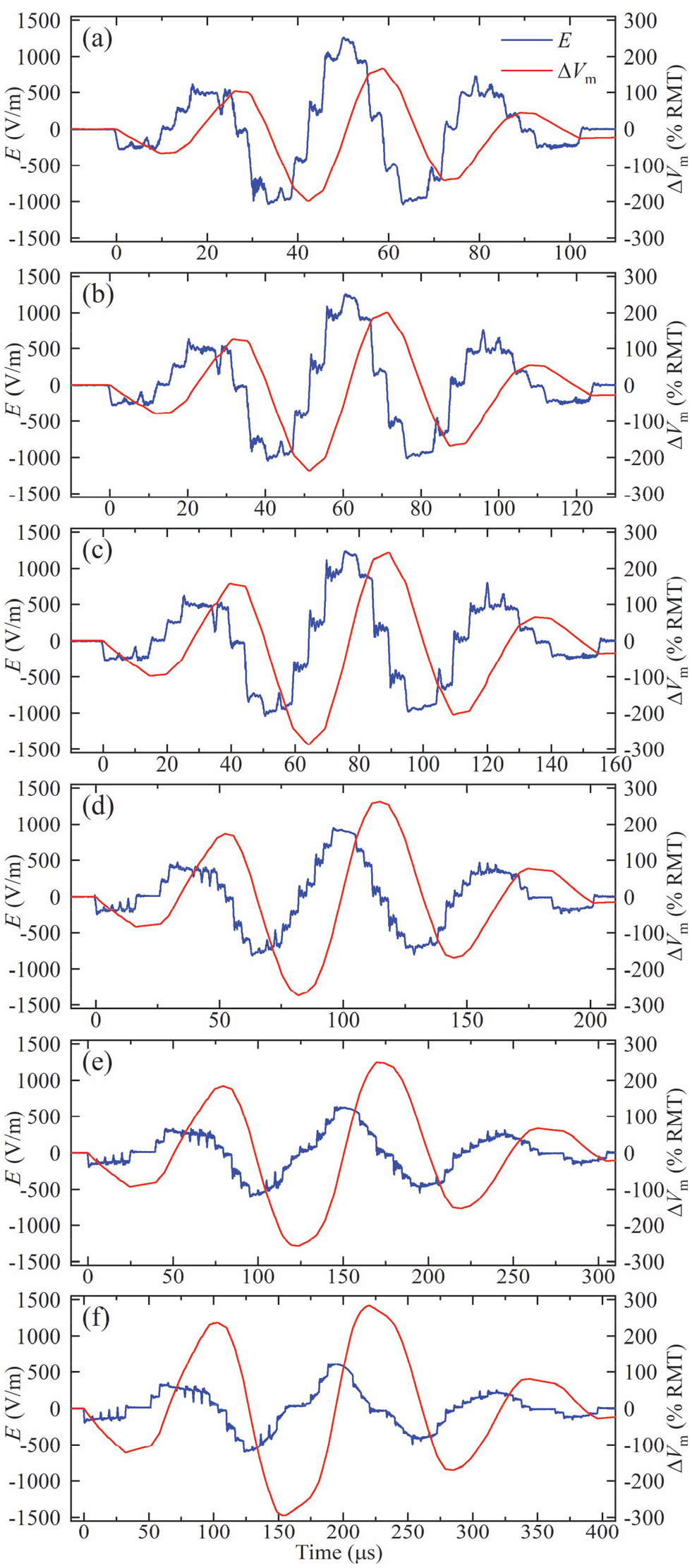
Measured electric field *E* and estimated neural membrane depolarization Δ*V*_m_ of the polyphasic pulses with Gaussian amplitude envelope with fundamental frequency of approximately (a) 30 kHz, (b) 25 kHz, (c) 20 kHz, (d) 15 kHz, (e) 10 kHz, and (f) 7.5 kHz. The initial module voltage *V*_*C*ref_ was selected so that the coil current during commutation did not exceed 6 kA.

### 4.6. Coil sound reduction

Since the sound emitted by a TMS coil is driven by electromagnetic forces, figure 11 shows spectral plots of the coil current and its square for representative MM-TMS pulses. In typical TMS applications, where there is no external magnetic field, the Lorentz forces within the coil are proportional to the cross product of the magnetic field and the current density, which is proportional to the square of the coil current. In specialized applications of TMS in the strong magnetic field of an MRI scanner, there are also Lorentz forces directly proportional to the coil current [53]. As expected, the coil current pulses have a spectral peak at the dominant pulse frequency, whereas the current squared displays peaks at twice that frequency.

**Figure 11.**
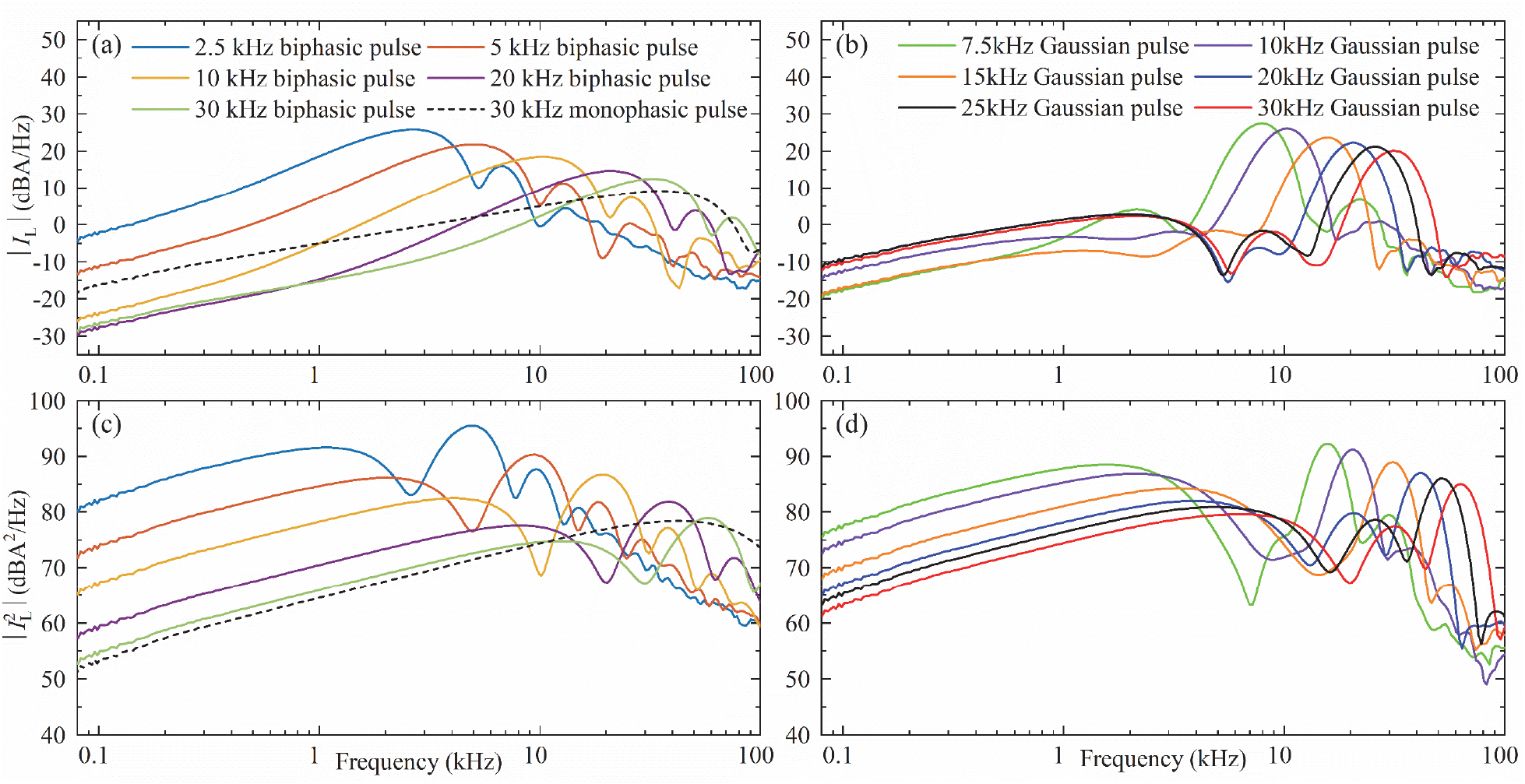
Smoothed 1/24-octave spectra of (a), (b) the measured coil current and (c), (d) the square of the coil current of selected biphasic rectangular pulses and the briefest monophasic rectangular pulse (left) as well as the polyphasic sinusoidal pulses with Gaussian amplitude envelope (right). The stimulation strength of each pulse was normalized to 167% average RMT.

Figures 12 and 13 as well as supplementary figures S13 and S14 show results of the acoustic recordings of the coil sound, which had the expected dependency on pulse duration. Figure 13 summarizes the pulse loudness for the various MM-TMS pules types illustrated in this paper. For matched estimated membrane depolarization based on the simplified linear first-order approximation—which has to be used with caution—briefer pulses were less loud than longer pulses. Further, the monophasic pulses were less loud than the biphasic pulses, which, in turn, were less loud than the Gaussian polyphasic pulses. The difference between monophasic and biphasic pulses was larger for briefer pulses, as, predictably, the biphasic pulses lost their depolarization efficiency advantage compared to monophasic pulses. Extrapolated to 5 cm from the coil surface, the DCC* coil with the briefest 17 µs monophasic pulse at 167% RMT had a peak sound pressure level (SPL) of 80 dB(Z) which is 32–55 dB lower than that of commercial TMS coils [2] and 14 dB lower than that of the DCC* coil with a conventional biphasic TMS pulse [40]. The continuous sound level (SL) for a simulated 20 Hz rTMS train was 64 dB(A), which is 26–45 dB less than that of commercial coils and 13 dB less than that of the DCC* coil with a conventional TMS pulse.

**Figure 12.**
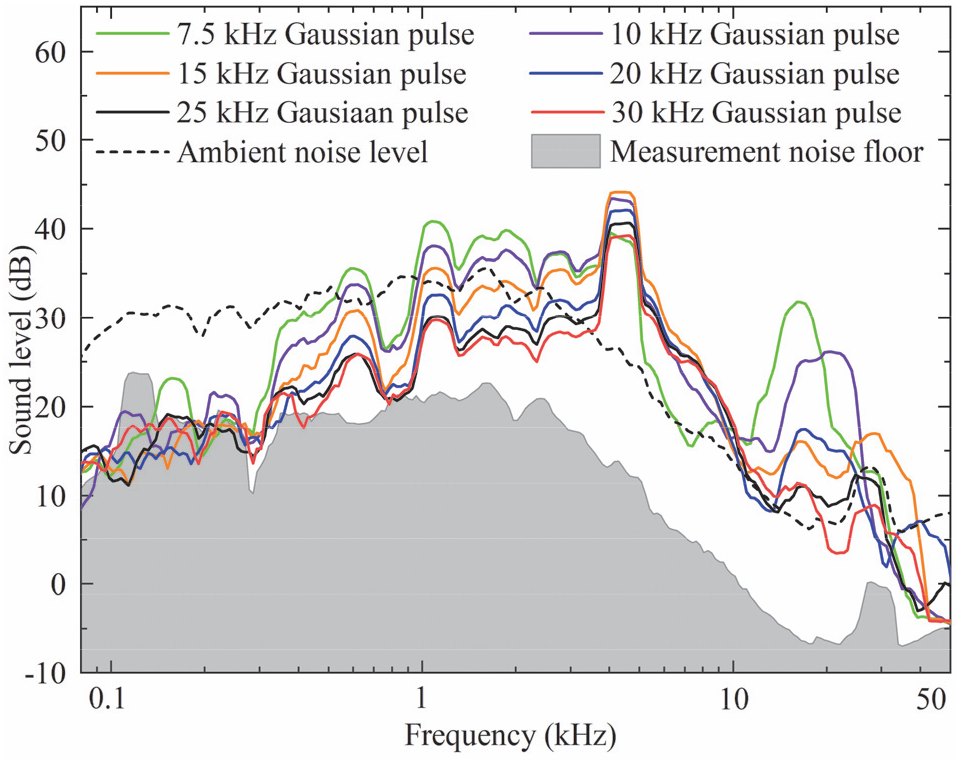
Smoothed 1/24-octave sound spectra of selected polyphasic sinusoidal pulses with Gaussian amplitude envelope. The sound has notable coil-specific components at 600, 1100, 2000, 2600, and 4300 Hz, and a pulse-specific component at twice the characteristic frequency of each pulse (15, 20, 30, and 40 kHz, respectively; the peaks for the two highest frequency pulses, 50 and 60 kHz, were above the recording bandwidth). The level of coil-specific components depends on pulse duration, and is in general lower for briefer pulses.

**Figure 13.**
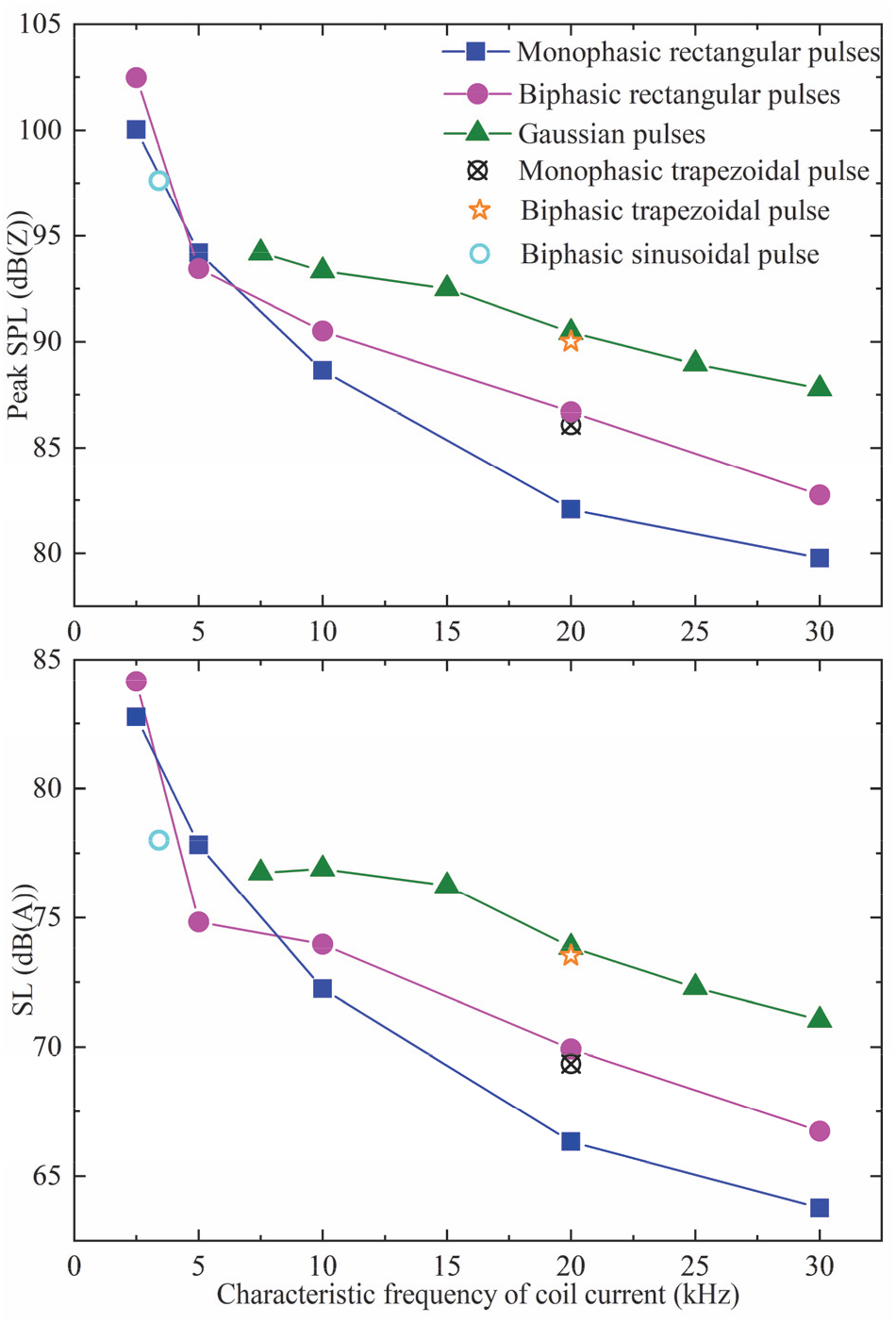
Peak sound pressure level (SPL) and sound level (SL) for various pulses generated by MM-TMS with the DCC* coil. SL is given for a simulated 20 Hz rTMS train and both SPL and SL are extrapolated to 5 cm from the coil surface for pulse amplitude of 167% average RMT.

The sound spectra of the different pulse waveforms show two mechanisms that explain this reduction. The Gaussian pulse acoustic spectra in figure 12 clearly show that the characteristic sound frequency of a TMS pulse is twice the characteristic electric field frequency. This is expected, as the Lorentz force is proportional to the coil current squared. Consequently, both briefer pulses and high-frequency Gaussian pulses push this part of the excitation forces and the sound energy out of the human hearing range, which is below 20 kHz. The attenuation is further amplified by the DCC* coil which was designed to work as an acoustic low-pass filter [40]. Further, for pulses with characteristic frequency much greater than the dominant modes of the DCC* (namely, the long and short modes of the winding block at around 2 and 4 kHz, respectively), briefer pulses have less subharmonic frequency content, reducing the sound intensity of these modes. This causes brief monophasic pulses to be quieter than their biphasic or Gaussian counterparts with matched frequency content (see supplementary figures S13 and S14 for the monophasic and biphasic spectra, respectively).

The sound differences between the pulses can be readily explained by differences in their squared-coil-current spectra (figure 11(c), (d)). The difference between the sound spectrum and the squared-coil-current spectrum is approximately a frequency-dependent constant factor—the acoustic transfer function of the coil. Using leave-one-out analysis on the pairs of sound and squared-current spectra, the geometric mean of their computed transfer functions can predict the left out sound spectrum from the corresponding squared-current spectrum with a mean prediction error of 2.4 dB and 95% percentile prediction error of 5.2 dB between 1 and 40 kHz. Below 1 kHz and above 40 kHz, the prediction accuracy deteriorates as the sound spectra hit the measurement noise floor. Notably, the 5 kHz biphasic pulse and the 7.5 kHz Gaussian envelope pulse were quieter than expected from the general trend for pulse durations in figure 13. This appears to be due to these two pulses having a minimum in their squared-coil-current spectra near the main sound-producing resonant frequency of the DCC* coil at around 4–5 kHz. It might be possible to further deepen this minimum and to tune it more accurately to the relatively narrowband resonant mode of the coil. Alternatively, it might be possible to generate a similar minimum for a briefer pulse with overall reduction in the squared-current spectrum in the hearing range.

## 5. Discussion

This paper presented a novel MM-TMS device with a wide and flexibly controlled output range of suprathreshold pulses. The MM-TMS device was able to generate the briefest TMS pulses reported to date, with biphasic duration as short as 33 μs. These high-voltage ultrabrief pulses had estimated neural membrane depolarization exceeding 300% RMT, suggesting suprathreshold stimulation strength. However, as noted earlier, the neural activation model used in this work has to be interpreted only as a rough approximation for the effective stimulation strength, since it simplifies significantly the neural dynamics and may underestimate the thresholds for ultrabrief pulses [45, 46]. In the future, these stimulation strength estimates can be replaced by empirical motor thresholds or more complex neural response models.

The MM-TMS device can generate a wider range of pulse shapes and widths (e.g., 33–400 μs biphasic) than previously possible [8-11, 27]. The ten-module MM-TMS device allows 21 output voltage levels in a pulse, which enables the generation of near-sinusoidal, near-rectangular, and more complex pulse shapes, not available in other TMS devices. We provided examples of complex polyphasic pulses with a Gaussian amplitude envelope and a wide range of pulse widths (99–400 μs). We had hypothesized that such pulses may reduce the pulse sound [22]. Our sound measurements in combination with the simplified linear threshold model suggest that the Gaussian envelope pulses with the specific carrier and envelope parameters selected here are louder than both the biphasic and monophasic pulses with the same characteristic frequency. The relative loudness of the Gaussian envelope pulses is due to a significant low-frequency sideband of the squared coil current, which is the dominant driving factor for the coil sound. However, these comparisons may be confounded since the simplified neural depolarization model is not able to correctly estimate the threshold for polyphasic pulses [28, 29, 47, 54]. Moreover, we did not optimize any parameters of the Gaussian pulses such as the width of the envelope. Worth noting is that the Gaussian envelope pulses are likely quieter than matched polyhasic pulses with a flat envelope [28, 29], although this was not tested expeirmentally. Finally, our measurements suggest that the coil sound reduction could be optimized by coordinating the spectra of the driving electromagnetic forces and the coil acoustic transfer function, for example by matching peaks in one to minima in the other.

The MM-TMS device implements efficient energy recycling since most of the pulse energy is recovered back to the capacitors. This, combined with the unprecedented flexible control over the pulse shape, enables a single device to generate rapid pulses sequences with different shapes for each pulse, which conventionally require two or more TMS devices to be combined. Importantly, the pulse amplitude of each pulse can be adjusted up or down independently of the amplitude of the prior pulse, which is critical for paired-pulse protocols.

Notably, except for the 1+1+1+1+1+1+1+1+1+1 sequential switching scheme intended for very long pulses, the MM-TMS device drives differentially the two coil terminals with a pair of voltages of equal magnitude but opposite polarity, leading to a zero common-mode voltage of the coil and zero voltage at the center of the conventional figure-of-8 TMS coil configuration at any instant during a pulse [26]. This could, in principle, reduce the artifact in concurrent EEG and EMG recordings resulting from high-voltage capacitive coupling between the TMS coil and the subject’s body and recording electronics.

Finally, a limitation of the MM-TMS circuit topology is that it allows only series or bypass connections among the modules. Thus, the total capacitance across the modules cannot be utilized at low output voltage levels since the modules cannot be connected in parallel. The MM-TMS topology can, however, be modified to add parallel connectivity [37]. While this is a compelling direction for future research, we did not pursue it in this work due to the added complexity of implementing and controlling the required additional IGBT switches.

## 6. Conclusion

We developed the first suprathreshold TMS device using a modular multilevel circuit topology at full TMS energy levels. The MM-TMS device allows unprecedented control of the pulse shape, amplitude, and width, which could enable improved and novel research, diagnostic, and therapeutic protocols, including ones with reduced acoustic noise.

## Acknowledgments

Research reported in this publication was supported by the National Institute of Mental Health of the National Institutes of Health under award numbers R01MH111865 and RF1MH124943 as well as by hardware donations from Magstim. The content is solely the responsibility of the authors and does not necessarily represent the official views of the National Institutes of Health.

L. M. Koponen, S. M. Goetz, and A. V. Peterchev are inventors on patents and patent applications on TMS technology, including the coil technology described in this paper. S. M. Goetz is also inventor on patents covering aspects of the TMS pulse generator. Related to magnetic stimulation technology, S. M. Goetz has received research funding and patent application support from Magstim, and A. V. Peterchev has received research funding, travel support, patent royalties, consulting fees, equipment loans, hardware donations, and/or patent application support from Rogue Research, Tal Medical/Neurex, Magstim, MagVenture, Neuronetics, BTL Industries, and Advise Connect Inspire.

