## Supplementary Material for "Modular multilevel TMS device with wide output range and ultrabrief pulse capability for sound reduction"

**Zhiyong Zeng[1], Lari M. Koponen[1], Rena Hamdan[1], Zhongxi Li[2], Stefan M. Goetz[1,2,3,4,*] and Angel V. Peterchev[1,2,3,4,5,*]**

[1] Department of Psychiatry and Behavioral Sciences, Duke University, NC 27710, USA
[2] Department of Electrical and Computer Engineering, Duke University, NC 27708, USA
[3] Duke Institute for Brain Sciences, Duke University, NC 27708, USA
[4] Department of Neurosurgery, Duke University, NC 27710, USA
[5] Department of Biomedical Engineering, Duke University, NC 27708, USA
[*] contributed equally

#### Active snubbing

##### *Brief pulse snubbing*

The snubbers, the stimulation coil, and the circuit parasitic resistance form an L-R-C resonant circuit after a TMS pulse, introducing undesirable electric field oscillation [1, 2]. For example, near the end of a negative biphasic pulse, the coil current flows through $D_{i4} \rightarrow C_i \rightarrow D_{i2}$, where $C_{i11}$, $C_{i12}$, $C_{i31}$, and $C_{i32}$ are charged to $V_{Ci}$ while $C_{i21}$, $C_{i22}$, $C_{i41}$, and $C_{i42}$ have a near-zero voltage. When the coil current decays to zero, each module outputs $-V_{Ci}$, and a high voltage $V_L = -10V_{Ci}$ is imposed on the stimulation coil, initiating a negative coil current. The negative current discharges capacitors $C_{i11}$, $C_{i12}$, $C_{i31}$, and $C_{i32}$ and charges capacitors $C_{i21}$, $C_{i22}$, $C_{i41}$, and $C_{i42}$. The voltage and current in the snubber capacitors and coil oscillate until the circuit resistance dissipates all the energy, which causes large artifacts at the end of the electric field pulses.

To dampen the ringing at the end of the pulse, the module can be switched between states with a high and low output voltage with a high frequency, which produces a controllable equivalent output resistance determined by the switching frequency and duty ratio [1]. The energy loss mechanisms resulting in the effective resistance are the increased IGBT collector–emitter resistance during switching and the resistive charge or discharge through the IGBT of the energy stored on the snubber capacitors. In the above example, switching $Q_{i2}$ rapidly between on and off state can provide a controllable higher equivalent output resistance. Moreover, since diode $D_{i2}$ is on during the last phase of the pulse ending in a positive current, the gate signal of $Q_{i2}$ does affect the TMS pulse. Therefore, the module state sequences for subbing can be activated before the end of the TMS pulse, and the active coil snubbing starts automatically when the coil current decays to zero [1]. Figure S1 diagrams this active snubbing scheme applied synchronously across all ten modules. Adjusting the snubber switching frequency, $T_s$, and duty ratio, $d = T_{on}/T_s$, controls the equivalent output resistance and therefore the damping. The snubbing action continues for $nT_s$, which should be chosen to extend sufficiently beyond the duration of the last phase of the pulse to suppress the coil current ringing. Figure S2 shows experimental data for this snubbing scheme and various duty ratios. While duty ratios of 0.6–0.7 supress rapidly the residual coil current, there is significant spiking in the electric field waveform.

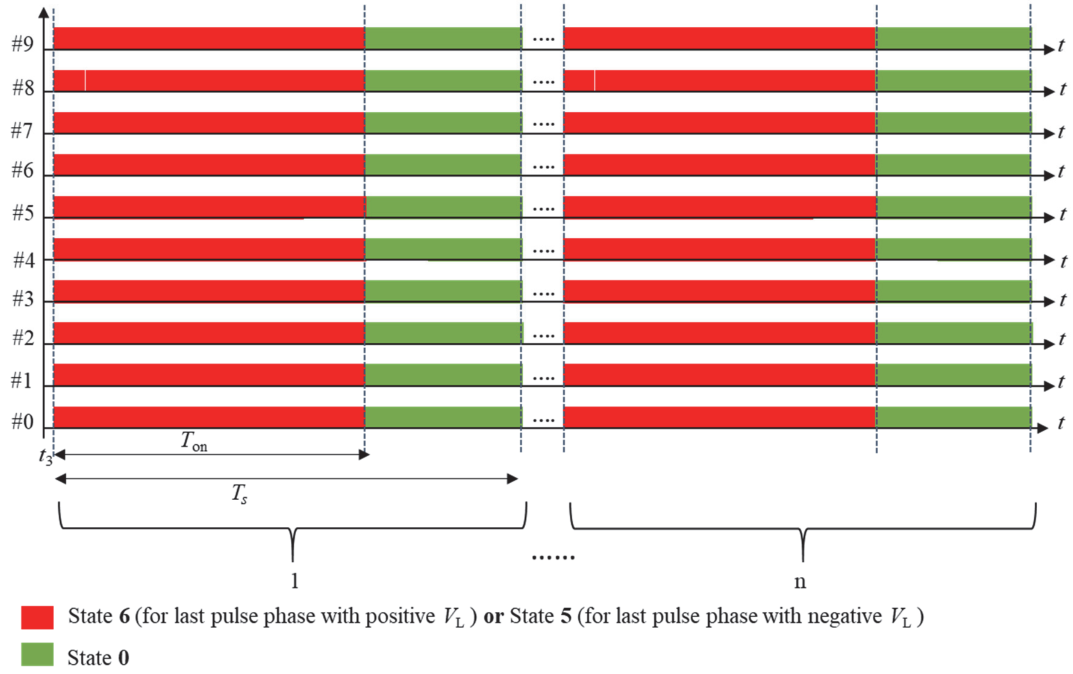


**Figure S1.** Module state timing diagram for synchronous active snubbing of a high-voltage brief TMS pulse. The module state numbers refer to table 1 in the main text. Time point $t_3$ denotes the beginning of the active snubbing, which lags behind the end of the pulse's second-to-last phase by a small interval to accommodate the switching transient. The controller repeats $n$ snubbing cycles of duration $T_s$. The module states before $t_3$ are not illustrated and can be found in the main text.

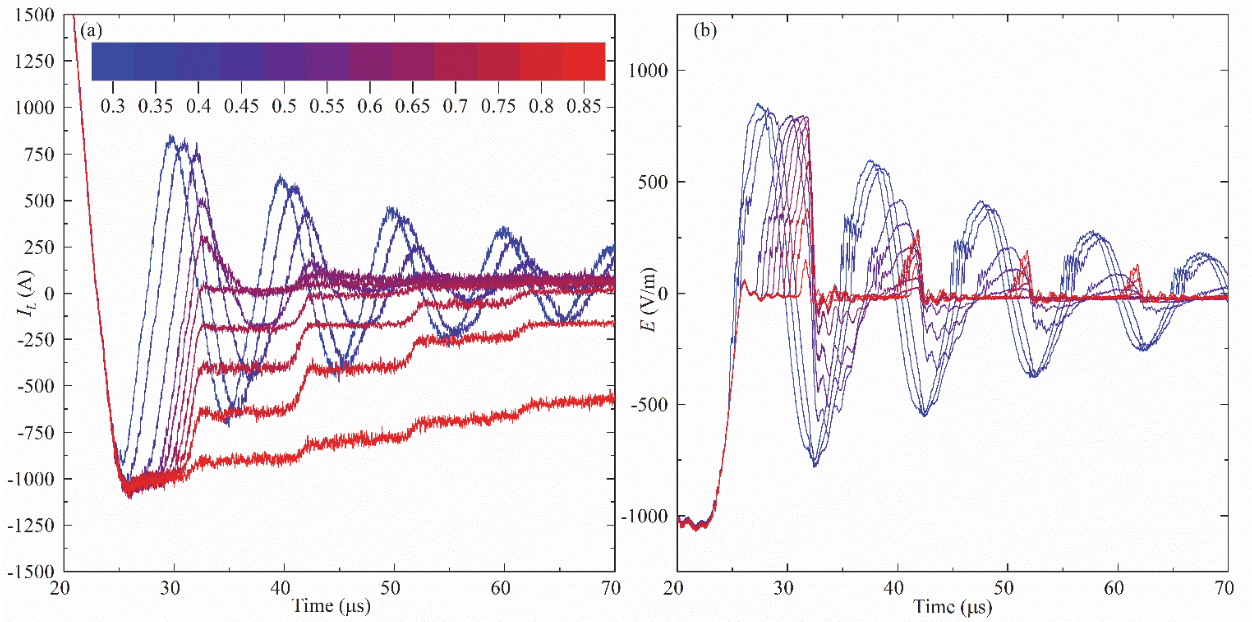


**Figure S2** Measurements of the synchronized snubbing scheme (figure S1) with duty ratio $d$ ranging from 0.3 to 0.85 at the end of a positive monophasic TMS pulse with peak coil current of $I_L = 8$ kA. (a) Coil current $I_L$. (b) Electric field $E$. The snubber switching frequency is $1/T_s = 100$ kHz, which is the maximum rating of the IGBT gate driver.

To reduce the switching ripple and enhance the damping associated with the active snubbing, the switching of the ten cascaded MM-TMS modules can be interleaved to increase the effective switching frequency by a factor of ten. To achieve this, the switching of the ten modules is phase shifted by $\delta = T_s/10$ (see figure S3), increasing tenfold the effective switching frequency seen by the coil. The interleaved snubbing significantly improves the damping performance compared to the synchronized snubbing scheme, as illustrated by the experimental results in figure S4. The damping waveforms are smoother than those in figure S2, and faster damping is achieved especially at low duty ratios.

Figure S4 indicates that duty ratios ranging from 0.5 to 0.65 provide a good performance balance, as they result in electric field spikes not exceeding one-fourth of the TMS pulse's peak electric field, while producing a fast current decay that leads to the reduced energy dissipation in the coil.

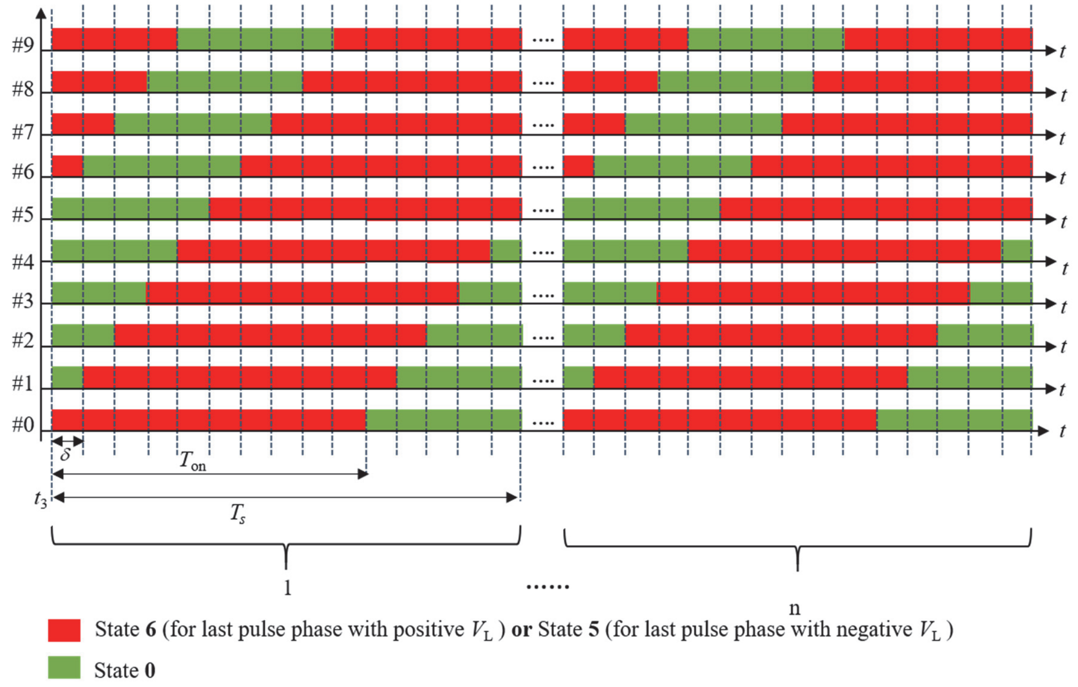


**Figure S3.** Module state timing diagram for interleaved active snubbing of a high-voltage brief TMS pulse. The modules' switching is interleaved by a phase shift of $\delta = T_s/10$. The conventions are the same as in figure S1.

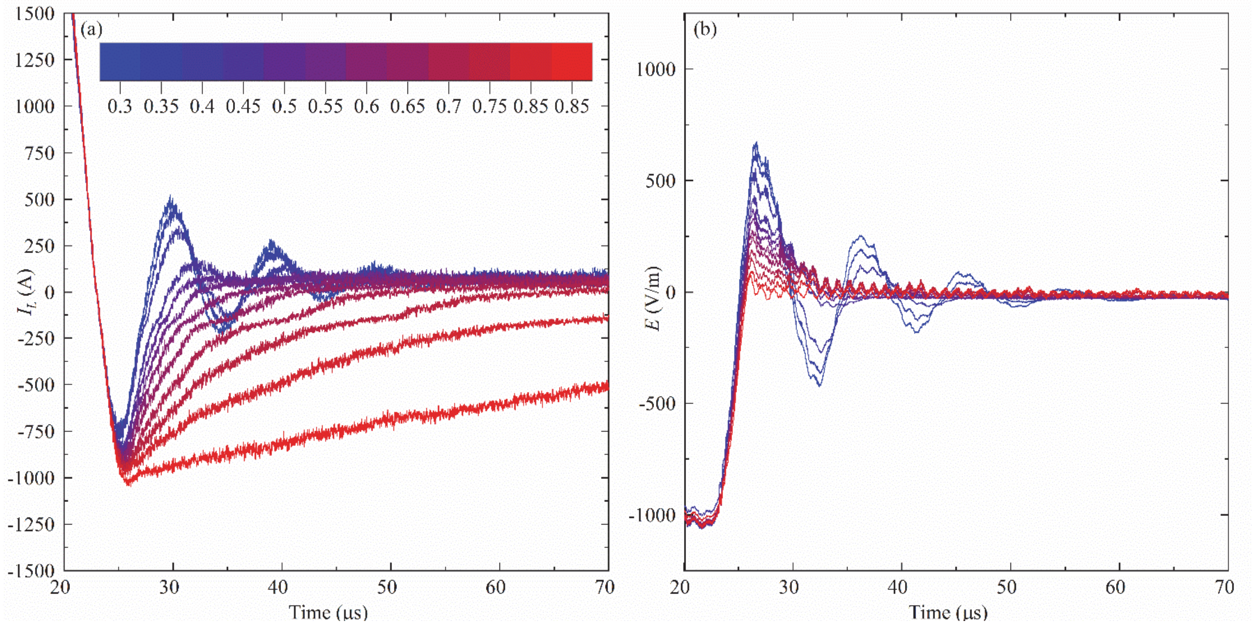


**Figure S4.** Measurements of the interleaved snubbing scheme (figure S3) with duty ratio $d$ ranging from 0.3 to 0.85 at the end of a positive monophasic TMS pulse with peak coil current of $I_L = 8$ kA. (a) Coil current $I_L$. (b) Electric field $E$. The snubber switching frequency is $1/T_s = 100$ kHz, which is the maximum rating of the IGBT gate driver.

*Long pulse snubbing*

Figures S5–S7 illustrate the module state timing for active snubbing of the long duration pulses shown in figure 6(g)–(l) in the main text. For these pulses, the MM-TMS modules are activated in sequential groups of 1+1+1+1+1+1+1+1+1+1, 2+2+2+2+2 and 5+5, respectively. The principles for designing the snubbing sequences were to wait for 1–2 $T_s$ cycles to ensure that the transition to the voltage transition to the last phase is complete, and then apply a switching pattern like in figure S1, but only the modules that are not in bypass mode. After the end of the last phase, these switching patterns apply brief damping pulses to the coil with voltage amplitude not exceeding that of the last phase. This snubbing approach was also used for the polyphasic pulses in figure 9 in the main text.

The snubbing schemes in figures S5–S7 can be refined in various ways. For example, after snubbing cycle 8 and 3 in figure S5 and S6, respectively, the snubbing switching can be rotated sequentially through all of the modules to distribute more evenly the switching and energy dissipation stress. Moreover, by interleaving the snubbing switching across all modules analogous to figure S3, the snubbing ripple can be reduced similarly to the performance shown in figure S4. This requires, however, more complex programming of the module states which was not implemented in this prototype.

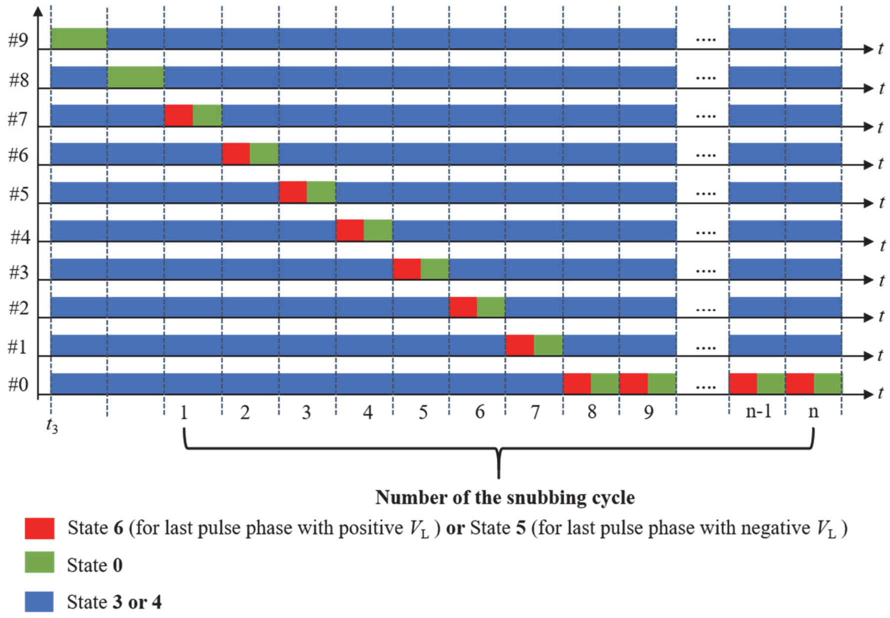


**Figure S5.** Module state timing diagram for active snubbing of a long duration TMS pulse using sequential module firing with the 1+1+1+1+1+1+1+1+1+1 scheme. The conventions are the same as in figure S1.

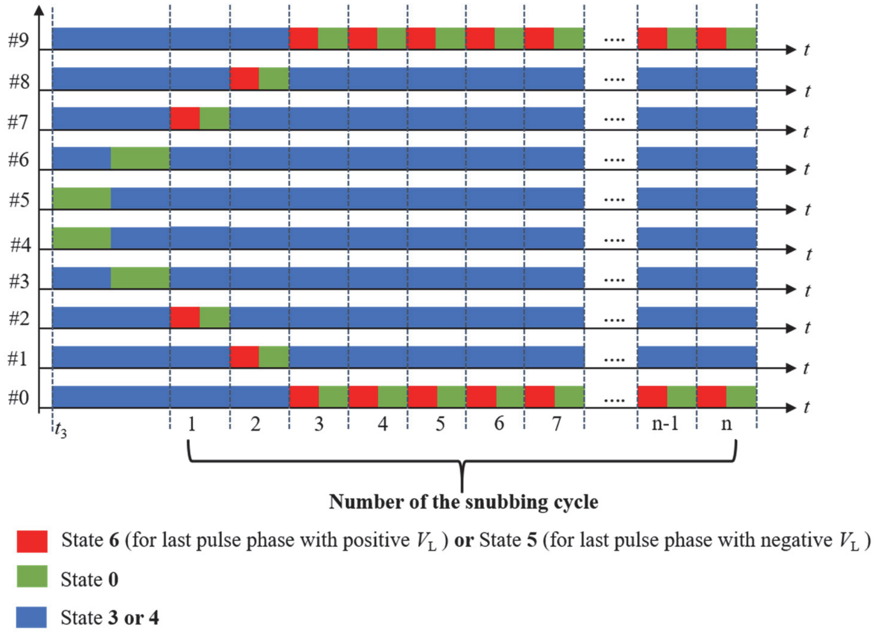

**Figure S6.** Module state timing diagram for active snubbing of a long duration TMS pulse using sequential module firing with the 2+2+2+2+2 scheme. The conventions are the same as in figure S1.

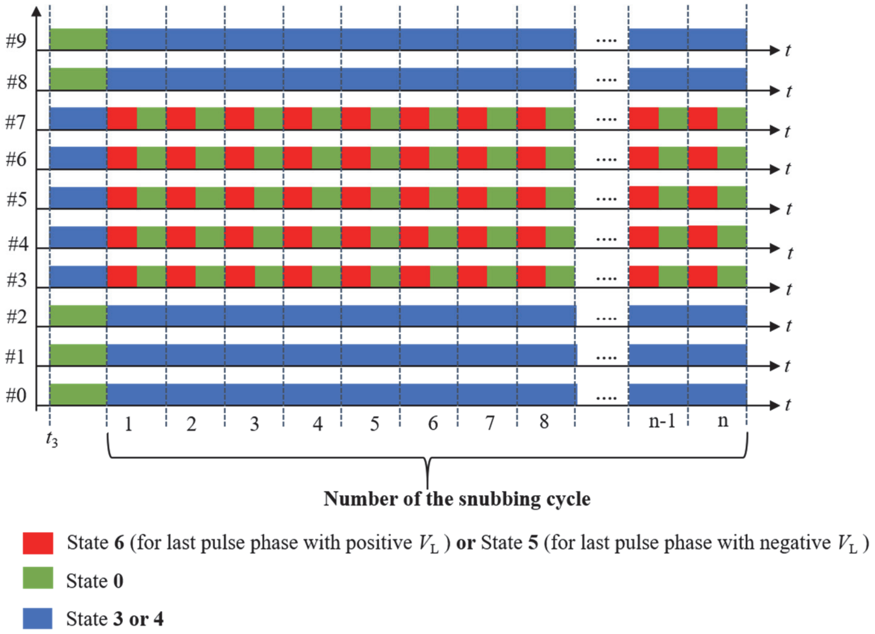


**Figure S7.** Module state timing diagram for active snubbing of a long duration TMS pulse using sequential module firing with the 5+5 modulation scheme. The conventions are the same as in figure S1.

### Extended duration pulses

Figures S8 and S9 show pulses that are are similar to those in figure 6(i)–(l), but have extended pulse width. This was achieved by extending the duration of the switching segments around the phase transitions from 10 µs to 30 µs. This was feasible, since there is no hard commutation of the coil current from diode to IGBT during these segments, and hence the maximum hard-commutation current during the pulses was not increased compared to those in figure 6(i)–(l).

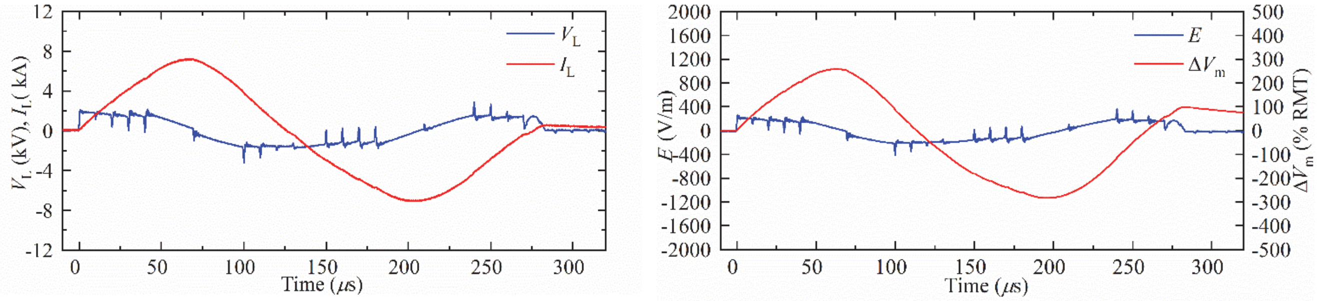


(a) (b)

**Figure S8.** Measured coil current $I_L$, coil voltage $V_L$, electric field $E$, and estimated neuro depolarization $\Delta V_m$ of positive biphasic pulses with a pulse width of 280 µs.

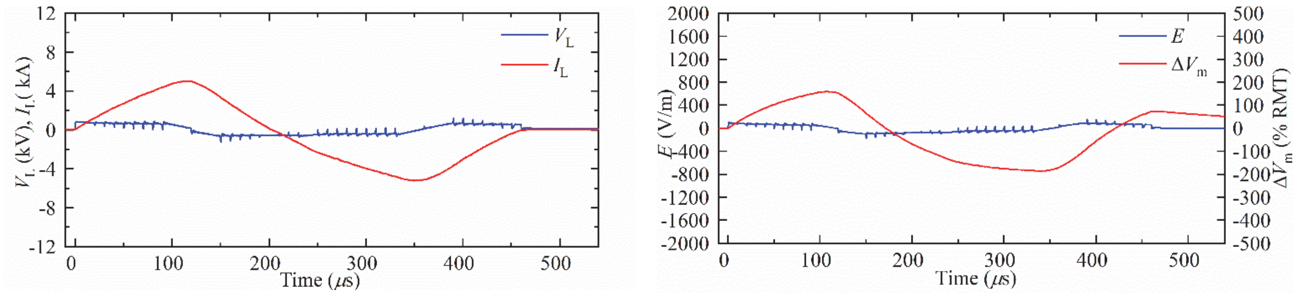


(a) (b)

**Figure S9.** Measured coil current $I_L$, coil voltage $V_L$, electric field $E$, and estimated neural depolarization $\Delta V_m$ of positive biphasic pulses with a pulse width of 480 µs.

### Pulse sequences

#### *Rapid sequential module activation*

Figure S10 shows a rapid sequence of positive biphasic TMS pulses generated by activating sequentially each of the ten MM-TMS modules with the remaining modules bypassed. The waveform demonstrates the ability to fire the modules is a rapid sequence as well as the nearly identical output of each module, which can be used as a diagnostic for correct functioning of the modules.

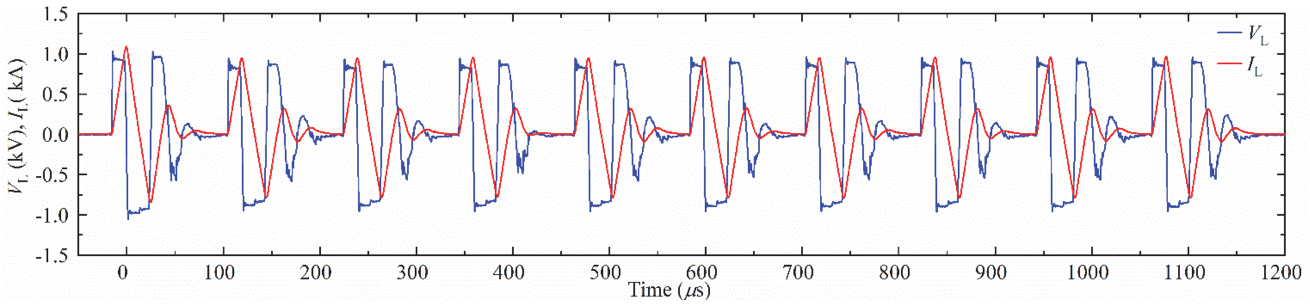


**Figure S10.** Measurements of a rapid train of positive biphasic pulses generated by activating each of the ten modules sequentially, while the remaining modules are bypassed.

#### *Paired pulses with different amplitudes*

Figure S11 shows pairs of TMS pulses identical to those in figure 10 in the main text but with a longer, 3 ms interstimulus interval.

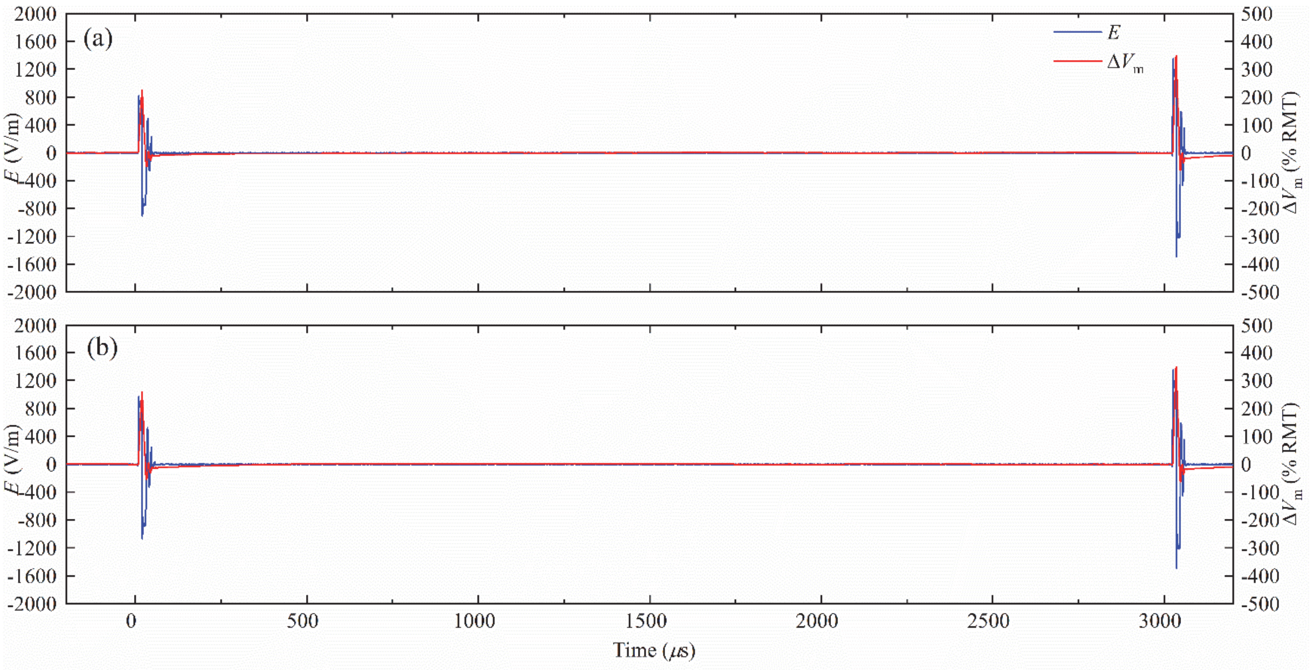


**Figure S11.** Pair of positive monophasic pulses with an interstimulus interval of 3 ms. (a) The first pulse has a lower stimulation strength than the second pulse by 36%. (b) The first pulse has a lower stimulation strength than the second pulse by 25%.

### Transient coil currents during module charging

As diagrammed in figure 1 in the main text, the charger's positive output is connected to the energy storage capacitor's positive terminal $p_{c0}$ of module #0, and the grounded charger's negative output is connected to the half-bridges output $n_0$ of module #0. When module #0 is charged, all IGBTs are off, and there is no transient current going through the stimulation coil when module #0 is charged. However, there is a transient current between module #0 and the target module when charging modules #1–#9. For example, when module #1 is charged, the unbalanced voltage between C1 and C31 introduces the transient current between module #0 and module #1 (see figure S12). The stimulation coil is a part of the conduction path for the inrush current. The inrush current is unavoidable with the proposed charging scheme. Nevertheless, the inrush current can be reduced by minimizing the voltage difference between modules by using several rounds of incremental charging of the modules.

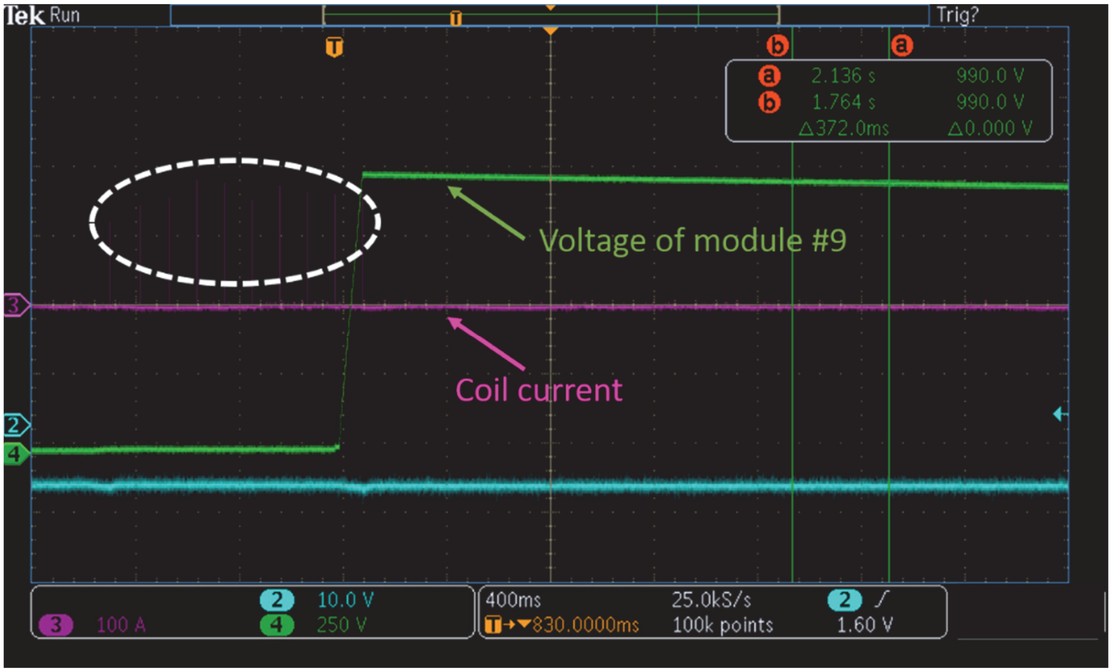


**Figure S12.** Transient coil current for initial charging when all modules have voltages close to zero.

### Pulse sound

Figures S13 and S14 shown spectral plots analogous to figure 12 in the main text but for monophasic and biphasic rectangular pulses respectively.

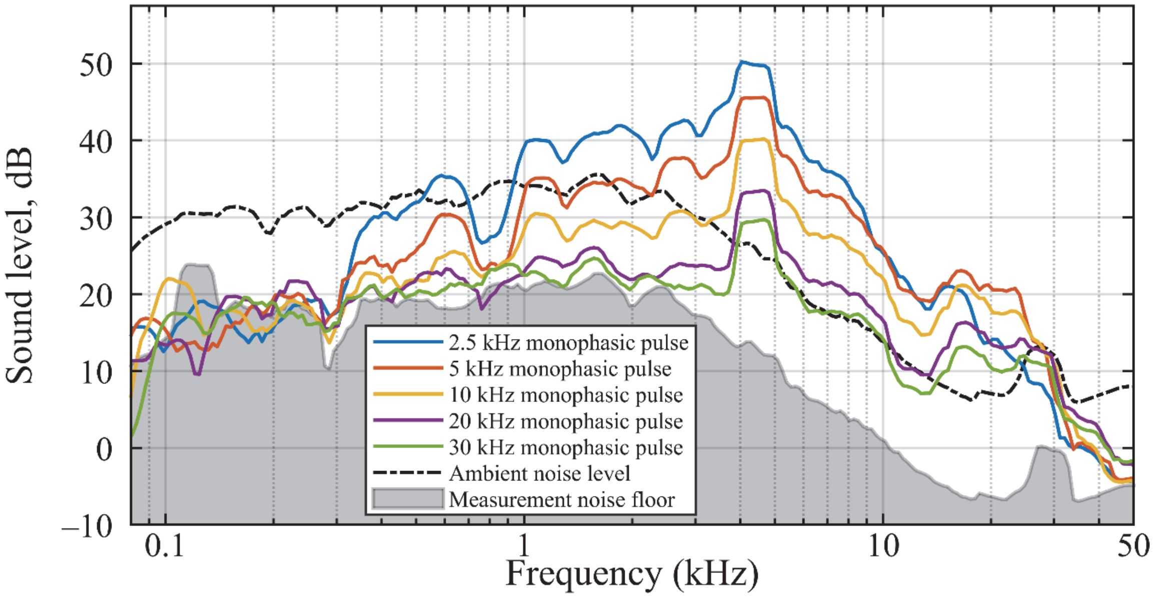


**Figure S13.** Smoothed 1/24-octave sound spectra of monophasic rectangular MM-TMS pulses. Reducing the pulse duration reduces the intensity of coil-specific low-frequency sound components. For rectangular monophasic pulses, a weak pulse-specific component can be seen at a frequency slightly below twice the nominal characteristic frequency of the pulse. Despite the averaging, similarly to figure 12, the sound spectra of two shortest pulses at frequencies below 2 kHz could not be reliably estimated due to very limited amount of low frequency content in these pulses.

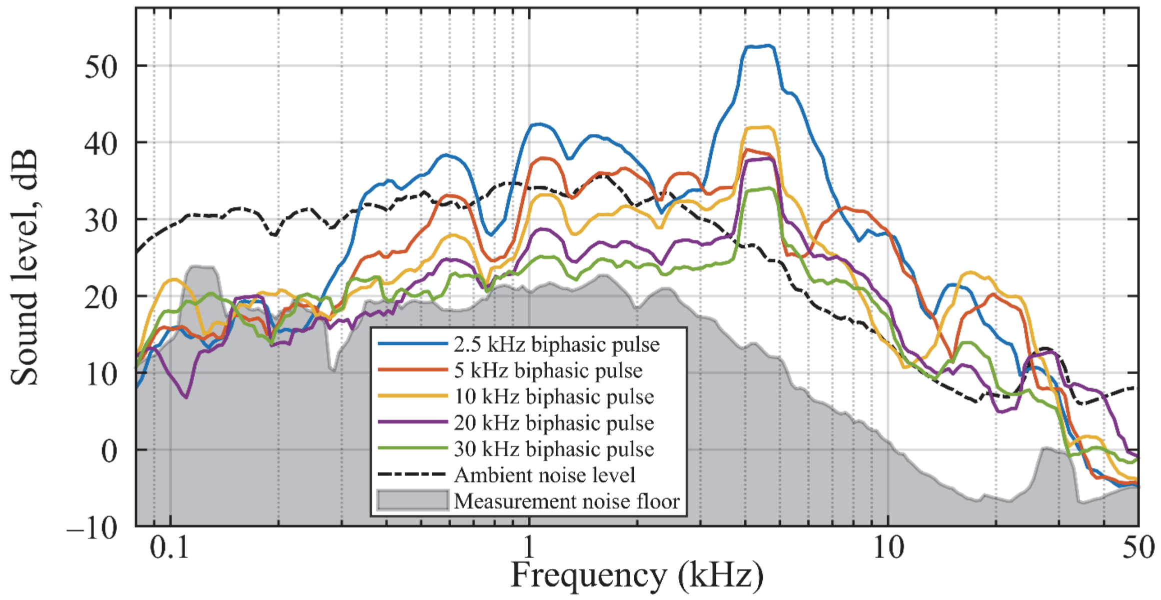


**Figure S14.** Smoothed 1/24-octave sound spectra of biphasic rectangular MM-TMS pulses. Reducing the pulse duration reduces the intensity of coil-specific low-frequency sound components. For rectangular biphasic pulses, a pulse-specific component can be seen at a frequency slightly below twice the nominal characteristic frequency of the pulse. For the longest pulse, this pulse-specific component couples with the coil-specific component at 4300 Hz, which increases the intensity of that component of sound, which can also manifest as an overall increase of both peak SPL and continuous SL of that pulse as shown in figure 13. An opposite phenomenon to this can be seen for the second longest pulse. In this case, the coil-specific component at 4300 Hz lines with the expected small reduction in pulse-specific intensity, which for biphasic pulses occurs at their characteristic frequency.
